# nAdder: A scale-space approach for the 3D analysis of neuronal traces

**DOI:** 10.1101/2020.06.01.127035

**Authors:** Minh Son Phan, Katherine Matho, Emmanuel Beaurepaire, Jean Livet, Anatole Chessel

## Abstract

Tridimensional microscopy and algorithms for automated segmentation and tracing are revolutionizing neuroscience through the generation of growing libraries of neuron reconstructions. Innovative computational methods are needed to analyze these neural traces. In particular, means to analyse the geometric properties of traced neurites along their trajectory have been lacking. Here, we propose a local tridimensional (3D) scale metric derived from differential geometry, which is the distance in micrometers along which a curve is fully 3D as opposed to being embedded in a 2D plane or 1D line. We apply this metric to various neuronal traces ranging from single neurons to whole brain data. By providing a local readout of the geometric complexity, it offers a new mean of describing and comparing axonal and dendritic arbors from individual neurons or the behavior of axonal projections in different brain regions. This broadly applicable approach termed nAdder is available through the GeNePy3D open-source Python quantitative geometry library.

## 1 Introduction

Throughout the history of the field of neuroscience, the analysis of single neuronal morphologies has played a major role in the classification of neuron types and the study of their function and development. The NeuroMorpho.Org database (Ascoli, 2006; Ascoli et al., 2007), which collects and indexes neuronal tracing data, currently hosts more than one hundred thousand arbors of diverse neurons from various animal species. Technological advances in large-scale electron (Helmstaedter et al., 2013; Kasthuri et al., 2015; Motta et al., 2019) and fluorescence microscopy (Gong et al., 2013; Winnubst et al., 2019; Abdeladim et al., 2019; Wang et al., 2019; Muñoz-Castañeda et al., 2020) facilitate the exploration of increasingly large volumes of brain tissue with ever improving resolution and contrast. These tridimensional (3D) imaging approaches are giving rise to a variety of model-centered trace sharing efforts such as the MouseLight (http://mouselight.janelia.org/, Winnubst et al., 2019), Zebrafish brain atlas (https://fishatlas.neuro.mpg.de/, Kunst et al., 2019) and drosophila connectome projects (https://neuprint.janelia.org/, Xu et al., 2020), among others.

Crucially, the coming of age of computer vision through advances in deep learning is offering ways to automate the extraction of neurite traces (Magliaro et al., 2019; Januszewski et al., 2018; Radojević & Meijering, 2019), a process both extremely tedious and time consuming when performed manually. This is currently resulting in a considerable increase in the amount of 3D neuron reconstructions from diverse species, brain regions, developmental stages and experimental conditions, holding the key to address multiple neuroscience questions (Meinertzhagen, 2018). Methods from quantitative and computational geometry will play a major role in handling and analyzing this growing body of neuronal reconstruction data in its full 3D complexity, a requisite to efficiently and accurately link the anatomy of neurons with varied aspects such as their function, development, and pathological or experimental alteration. Furthermore, morphological information is of crucial importance to address the issue of neuronal cell type classification, in complement to molecular data (Bates et al., 2019; Adkins et al., 2020).

An array of geometric algorithmic methods and associated software has already been developed to process neuronal reconstructions (Bates et al., 2020; Cuntz et al., 2010; Arshadi et al., 2020). Morphological features enabling the construction of neuron ontologies, described for instance by the Petilla convention (Petilla Interneuron Nomenclature Group et al., 2008), have been used for machine learning-based automated neuronal classification (Mihaljević et al., 2018). These features have also been exploited to address more targeted questions, such as comparing neuronal arbors across different experimental conditions in order to study the mechanisms controlling their geometry (Santos et al., 2018). In the context of large-scale traces across entire brain structures or the whole brain, morphological measurements are employed for an expanding range of purposes, e.g. to proofread reconstructions (Schneider-Mizell et al., 2016), probe changes in neuronal structure and connectivity during development (Gerhard et al., 2017) or identify new neuron subtypes (Wang et al., 2019).

So far however, metrics classically used to study neuronal traces tend to rely on elementary parameters such as length, direction and branching; as such, they do not enable to finely characterize and analyze neurite trajectories. One particularly interesting parameter that characterizes neurite traces is their local geometrical complexity, i.e. whether they adopt a straight or convoluted path at a given point along their trajectory. This parameter is of particular relevance for circuit studies, as it is tightly linked to axons’ and dendrites’ assembly and their role in information processing: indeed, axons typically follow simple paths within tracts while adopting a more complex structure at the level of terminal arbor branches that form synapses. Moreover, the sculpting of axonal arbors by branch elimination during circuit maturation can result in convoluted paths (Keller-Peck et al., 2001; Lu et al., 2009). The complexity of a neurite segment thus informs both on its function and developmental history. Available metrics such as tortuosity (the ratio of curvilinear to Euclidean distance along the path of a neurite) are generally global and average out local characteristics; moreover, traces from different neuronal types, brain regions or species can span vastly different volumes and exhibit curvature motifs over a variety of scales, making it difficult to choose which scale is most relevant for the analysis. One would therefore benefit from a generic method enabling to analyze the geometrical complexity of neuronal trajectories 1) locally, i.e. at each point of a trace, and 2) across a range of scales rather than at a single, arbitrary scale.

Methods based on differential geometry have been very successfully applied in pattern recognition and classic computer vision (Sapiro, 2006; Cao, 2003). In particular, the concept of scale-space has led to thorough theoretical developments and practical applications. The key idea is that starting from an original signal (an image, a curve, a time series, etc.), one can derive a family of signals that estimate that original signal as viewed at various spatial or temporal scales. This enables to select and focus the analysis on a specific scale of interest, or to remove noise or a low frequency background signal; in addition, scale-space analysis also provides multi-scale descriptions (Lindeberg, 1990). On 2D curves, the mean curvature motion is a well-defined and broadly applied scale-space computation algorithm (Cao, 2003). So far, however, comparatively little has been proposed concerning 3D curves, such data being less common (Digne et al., 2011). Applications of curvature and torsion scale-space analyses for 3D curves have been reported (Mokhtarian, 1988) but remain few and preliminary; the inherent mathematical difficulty represents another reason for this gap. Neuronal trace analysis clearly provides an incentive for further investigation of the issue.

Here, we present a complete framework for scale-space analysis of 3D curves and applied it to neuronal arbors. This method, which we name *nAdder* for Neurite Analysis through Dimensional Decomposition in Elementary Regions, allows us to compute the local 3D scale along a curve, which is the size in micrometers of its 3D structure, i.e. the size at which the curve locally requires the three dimensions of space to be described. This scale is quite small for a very straight trace, and larger when the trace displays a complex and convoluted pattern over longer distances. We propose examples and applications on several published neuronal trace datasets that demonstrate the interest of this metric to compare the arbors of individual neurons, extract morphological features reflecting local changes in neurite behavior, and provide a novel region-level descriptor to enrich whole brain analyses. Implementation of the nAdder algorithms along with an array of geometry routines and functions are made available in the recently published GeNePy3D Python library (Phan & Chessel, 2020), available at https://genepy3d.gitlab.io. Code to reproduce all figures in this study, exemplifying its use, is available at https://gitlab.com/msphan/multiscale-intrinsic-dim.

## 2 Results

We introduce a new form of analysis to investigate the spatial complexity of neuronal arbors. Our approach consists of decomposing each branch of a neuronal arbor into a sequence of curve fragments which are each assigned an intrinsic dimensionality, i.e. the smallest number of dimensions that accurately describes them, from which we compute a local 3D scale, defined as the highest scale at which the curve locally has a 3D intrinsic dimensionality. The first sub-section of the Results details the principle underlying this approach. We next demonstrate its purpose, accuracy and relevance through several applications.

### 2.1 Computation of the local 3D scale of neuronal traces from their multiscale intrinsic dimensionality

Given a neuronal arbor, we decompose each of its branches into a sequence of local curved fragments, which are classified according to their intrinsic dimensionality, defined by combining computations of both curvature and torsion of the curves. Briefly, a portion of curve with low torsion and curvature would be considered a 1D line, one with high curvature but low torsion would be approximately embedded within a 2D plane, and a curve with high torsion could only be described by taking into account the three spatial dimensions. This results in a hierarchical decomposition, capturing the fact that a 1D line is basically embedded in a 2D plane (Figure 1A, S1). To validate the proposed decomposition, we applied it on simulated traces presenting different noise levels. Our algorithm reached accuracies above 90% with respect to the known dimensionality of the simulations at low noise level and still above 80% at high noise level, while an approach not based on space-scale which we took as baseline (Yang et al., 2016; Ma et al., 2017) was strongly affected by noise (Figure 1A, S2 and S3; details on the simulation of traces, noise levels and algorithm are available in the Methods section).

**Figure 1:**
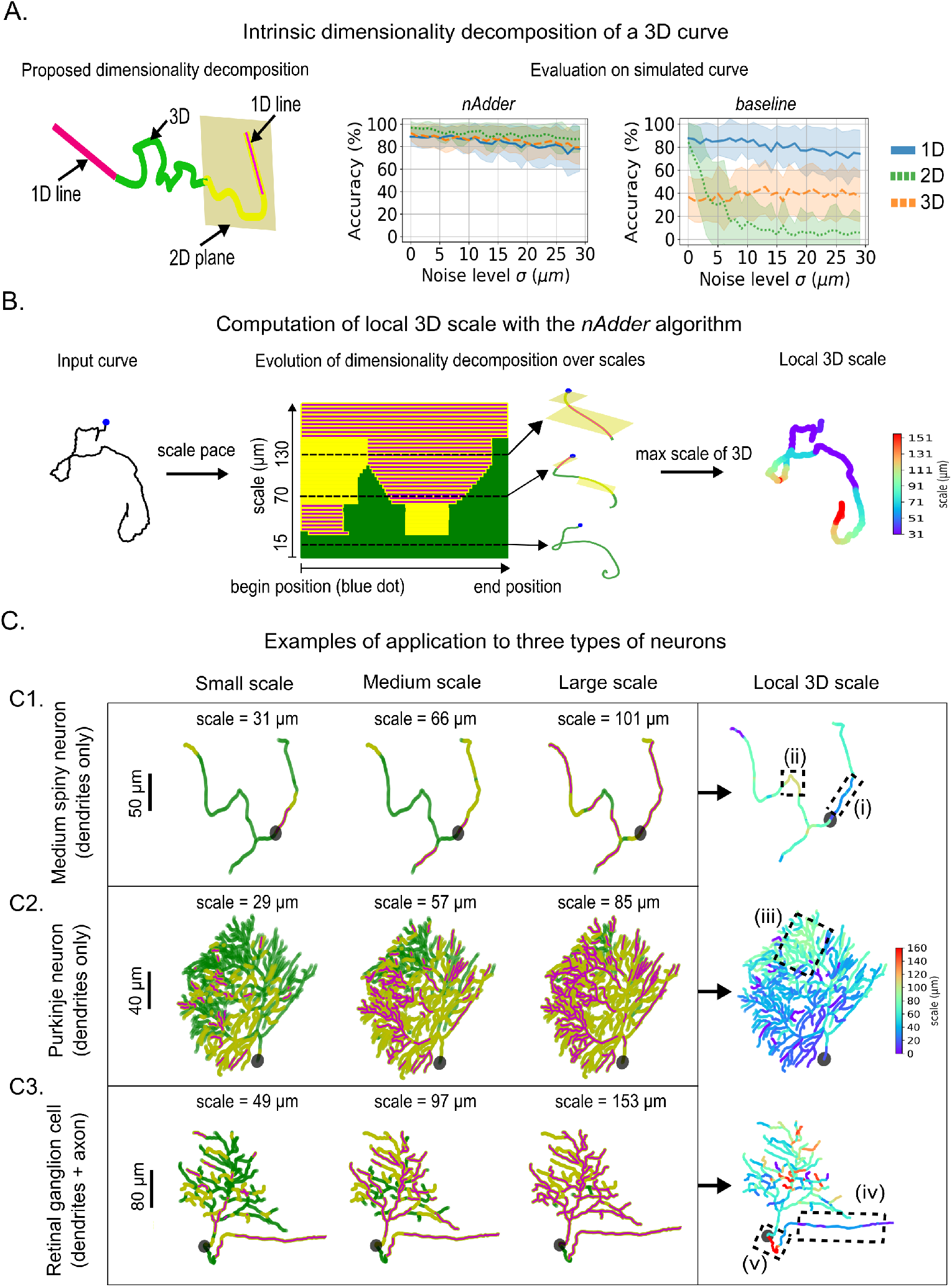
Computation of the 3D local scale of neuronal traces from their multiscale dimensionality decomposition. (A) (left) Variation of the intrinsic dimensionality along a portion of 3D curve; (right) evaluation of the decomposition on simulated trajectories and comparison to a baseline method not based on scale-space (Yang et al., 2016; Ma et al., 2017). (B) Schematic presentation of the processing of 3D curves by the nAdder algorithm. (C) Application of nAdder to three types of mouse neurons with different arbor shape and size: (C1) striatal D2 medium spiny neuron (Li et al., 2019), (C2) cerebellar Purkinje neuron (Chen et al., 2013), (C3) retinal ganglion cell (Badea & Nathans, 2011). The first three columns show the intrinsic dimensionality decompositions of the neurons at small, medium and large scales. The local 3D scale, computed from suites of such decomposition across multiple scales, is shown in the last column. The maximum scale is set at 160 *μ*m based on the longest branch analyzed. Dotted line boxes (i-v) frame areas of interest discussed in the text.

Our proposed decomposition of traces according to their local intrinsic dimensionality depends on the scale at which the analysis is performed. To develop a metric based on this decomposition that would be robust to scale, we implemented a space-scale approach. The notion of scale can be abstractly interpreted as the level of detail that an observer takes into account when considering an object, varying from a high level (when observing a trace up close i.e. at smaller scales), to low level (when observing it from afar i.e. at larger scales). A scale-space is the computation of all versions of a given curve across spatial scales. Several mathematical and computational frameworks have been proposed to compute such ensemble of curves; the simplest one used here consists of smoothing the initial portion of trace analyzed by convolutions of its coordinates with Gaussian kernels. The dimensionality of such a curve element, as we analyze it at increasing scales, is typically best described as 3D at the smallest scales (i.e. taking in account a high level of detail), and becomes 2D or 1D at larger scales (i.e. low level of detail) as more and more details are smoothed out (Figure 1B, middle panel and S4). Here, in the context of neuronal arbors, scales are measured in micrometers, by computing the radius of curvature of the smallest detail kept at that scale (see methods for details).

To make use of the local dimensionality decomposition across multiple scales and compute a simpler and more intuitive metric, we define the local 3D scale as the highest scale at which the trace still remains 3D. The trace will then locally transform to 2D or 1D for scales higher than that local 3D scale. An example of local 3D scale calculation is shown in Figure 1B. This computation is done at the level of curves; to apply it to whole neuronal arbors, we decompose the arbor into curves by considering, for each ‘leaf’ (terminals), the curves that link it to the root (i.e. the cell body), averaging values when needed (Figure S5). The result is an algorithm that computes a local 3D scale metric for full neuronal arbors, which we term *nAdder* (see Methods for more details on the algorithm).

To test our approach, we examined the local 3D scales of different types of neurons presenting dissimilar arbor shapes, using traces from the NeuroMorpho.Org database (Figure 1C). We first studied a mouse striatal D2-type medium spiny neuron reconstructed by Li et al. (2019) presenting a relatively simple structure with only a few dendrites (Figure 1C1, Movie S1). Computation of the local 3D scale for this neuron revealed variations in spatial structure across local regions. For instance, regions (i) and (ii) in Figure 1C1 correspond to two branch segments located at different distances from the soma, and the straightest segment (i) transforms from 3D to 2D and then 1D at a scale smaller than the more tortuous portion (ii). This results in a lower local 3D scale in region (i) (~40 *μ*m) compared to region (ii) (~100 *μ*m).

We then focused on a mouse cerebellar Purkinje neuron from Chen et al. (2013), presenting a more complex dendritic arbor (Figure 1C2, Movie S2). Most of this arbor was characterized by low local 3D scales (~45 *μ*m on average), in accordance with the fact that Purkinje cells dendritic arbors have a nearly flat appearance, and hence transform quickly to a 2D plane when scanning the scale space. Nevertheless, the analysis revealed one region (iii) presenting local 3D scales higher than the mean value, which was linked to the fact that dendrites in this region protruded in an unusual manner out of the main plane of the arbor (see Movie S2).

Finally, we examined the reconstruction of a mouse retinal ganglion cell (Badea & Nathans, 2011) comprising its entire dendritic arbor and the portion of its axon enclosed in the retina (Figure 1C3, Movie S3). We observed small local 3D scales (~35 *μ*m) along most of the axonal trajectory (see region (iv) in 1C3) except in its most proximal portion characterized by high local 3D scales (~150 *μ*m, region (v)). This reflects the structure of the axon as it first courses in a tortuous manner towards the inner side of the retina before running more directly towards the optic nerve head, the exit point for ganglion cell axons leaving the retina. The proximal region of the axon thus converts from 3D to 2D and 1D much faster than the more distal portion. The dendritic arbor presented an intermediate local 3D scale on the whole (~80 *μ*m) but with strong local variations. This corresponded to the fact that some dendritic branches exhibited complex shapes whereas others were more direct, as can be seen in Movies S2 and S3.

We also compared our local 3D scale metric with two classic local descriptors of 3D traces, curvature and torsion (Figure S6). Mapping of the three parameters in the Purkinje neuron presented in Figure 1C2 showed that the local 3D scale provided the most informative measure of the geometric complexity of neuronal arbors, whereas curvature and torsion yielded representations that were difficult to interpret as they were highly discontinuous because of local variations of the curves.

Overall, the local 3D scale metric computed with *nAdder* offers a new local measure of the geometric complexity of neuronal traces, and provides information robust across neuron types, shapes and sizes. We subsequently used our algorithm to reanalyze published datasets in order to explore its capacity to extract biologically meaningful anatomical insights.

### 2.2 Application of *nAdder* to characterize neuronal arbors during growth and in experimentally altered condition

To evaluate the relevance and usefulness of the local 3D scale measure, we first applied *nAdder* to compare neuronal arbors at different stages of their development and in normal vs. experimentally altered contexts. We selected data from Santos et al. (2018) hosted in the NeuroMorpho.Org database which describe the expansion of the dendritic arbors of the Xenopus laevis tectal neuron during development and the effect of altering the expression of Down syndrome cell adhesion molecule (DSCAM). In this study, the authors showed that downregulation of DSCAM in tectal neuron dendritic arbors increases the total dendritic length and number branches, while overexpression of DSCAM lowers them. We reanalyzed neuronal traces generated from the dendritic arbors in this study by computing their local 3D scale using *nAdder*.

Our analysis showed that the mean local 3D scale of developing X. laevis tectal neurons overexpressing DSCAM is smaller than that of control neurons, with a significant difference between the two conditions, 24 and 48 hr after the start of the observations at Stage 45 (Figure 2A and B); this result is consistent with the original analysis by Santos et al. (2018), indicating that DSCAM overexpression leads to more simple arbor morphologies, as measured by the number of branches and length of reconstructed arbors. In their paper, the authors also studied tortuosity, the ratio of curvilinear to Euclidean length of a trace, a global morphological metric classically used to measure geometric complexity. While the authors show that DSCAM downregulation increases the tortuosity of the neuron’s longest dendritic branch (see Figure 4 of Santos et al. (2018)), they did not report this parameter for DSCAM overexpression. We computed the tortuosity of the longest branch and indeed observed no significant difference between control and DSCAM overexpressing neurons at any stage of the analysis (Figure 2C, middle). Interestingly, however, we found that the mean local 3D scale of the arbors’ longest branch (Figure 2C, top) was significant reduced in DSCAM-overexpressing vs. control neurons 24 and 48 hr after the start of the image series. We also observed that the length and mean local 3D scale of the neuron’s longest arbor branch were correlated in control neurons (Figure 2D, left) (r = 0.30, p value = 0.020), but disappeared in DSCAM-overexpressing neurons (Figure 2D, right). This suggests a subtle effect of DSCAM in uncoupling the length of neurites and their geometric complexity. Our analysis shows that measuring the neurites’ local 3D scale provides biologically relevant information and can measure subtle differences in the complexity of their trajectory that are not apparent with classic metrics.

**Figure 2:**
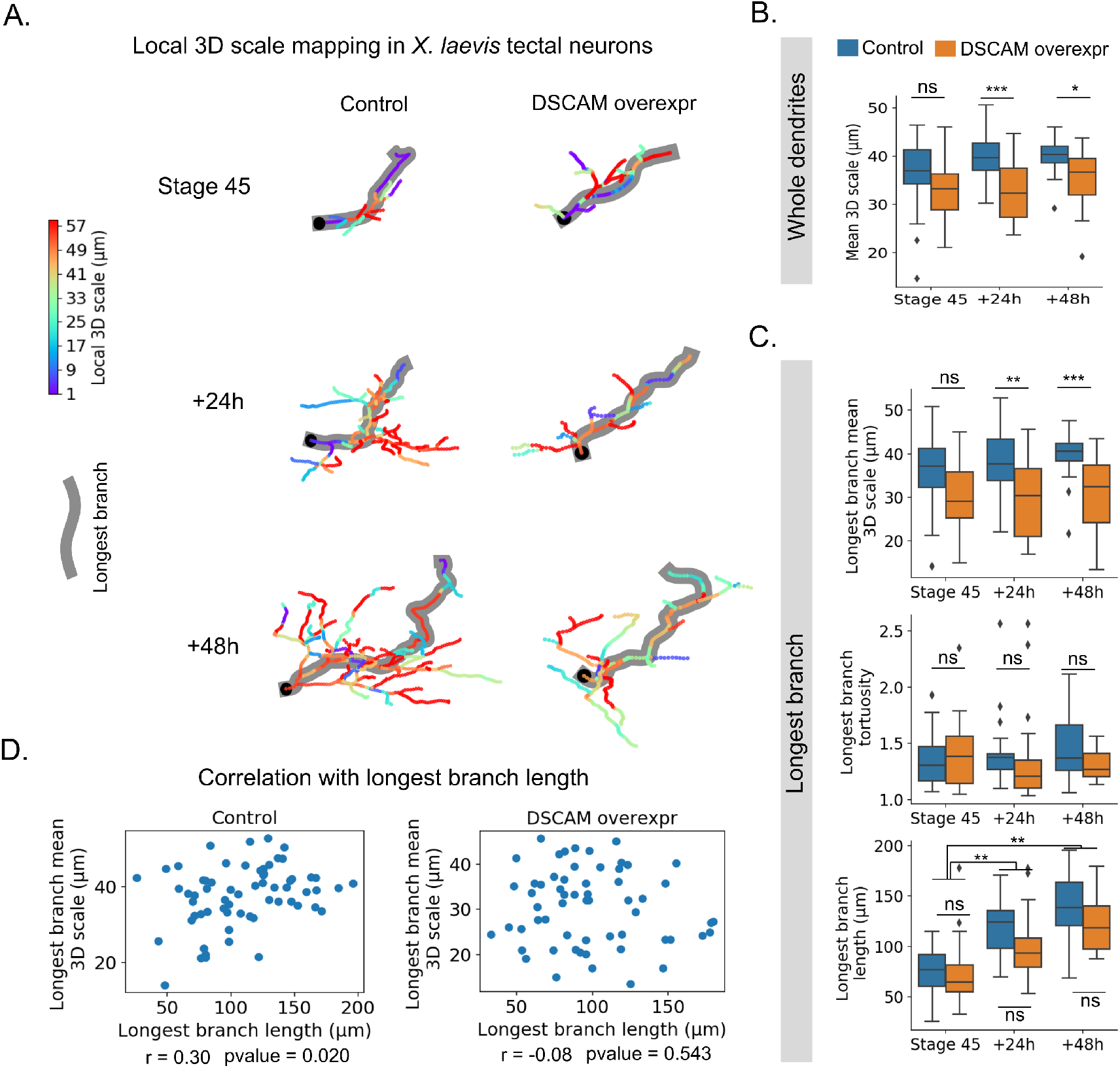
Local 3D scale analysis in tectal neurons from developing Xenopus laevis tadpoles and effect of DSCAM overexpression. Dendritic arbor traces from X. laevis tectal neurons were obtained from Santos et al. (2018). (A) Examples of local 3D scale maps from control (left) and DSCAM-overexpressing (DSCAM overexpr) neurons at Stage 45, and 24 and 48 hr after initial imaging. The maximum scale is set to 60 *μ*m based on the mean length of the arbor’s longest branch (highlighted in gray). The cell body’s position is indicated by a black square. (B) Evolution of the mean local 3D scale of the whole dendritic arbor from Stage 45, to +24 and +48 hr. (C) Mean local 3D scale (top), tortuosity (middle) and length (bottom) of the longest branch only. Two-way ANOVA, Student’s t-test with Holm-Sidak for multiple comparisons were used, * p≤0.05, ** p≤0.01, *** p≤0.005, **** p≤0.001. (D) Correlation between the mean 3D scale and the longest branch length in control vs. DSCAM-overexpression conditions. Spearman correlation was used.

### 2.3 Local 3D scale mapping across the whole larval zebrafish brain

We next sought to test *nAdder* over a large-scale 3D dataset encompassing entire long-range axonal projections. A database including 1939 individual neurons traced across larval zebrafish brains and co-registered within a shared framework has been published by Kunst et al. (2019). The authors clustered neuronal traces into distinct morphological classes using the NBLAST algorithm (Costa et al., 2016), thus identifying both known neuronal types and putative new ones. These neuronal traces were also used to build a connectivity diagram between 36×2 bilateral symmetric regions identified in the fish brain, revealing different connectivity patterns in the fore-, mid- and hindbrain. Here, we reanalyzed this dataset by computing the local 3D scales of all available neuronal traces, focusing on projections linking distinct brain regions (i.e. that terminated in a region distinct from that of the cell body), interpreted as axons. We then explored the resulting whole-brain map of the geometric complexity of projection neurons.

#### Local 3D scale variations across brain regions

Processing of the 1939 traces of the larval zebrafish atlas with *nAdder* resulted in a dataset where the local 3D scale of each inter-region projection (i.e. projecting across at least two distinct regions), when superimposed in the same referential, could be visualized (Figure 3A and S7A). Coordinated variations in local 3D scale values among neighboring traces were readily apparent on this map, such as in the retina (see region (i) in Figure 3A) and torus semicircularis (ii) which presented high local 3D scale values, or the octaval ganglion (iii) characterized by low values.

**Figure 3:**
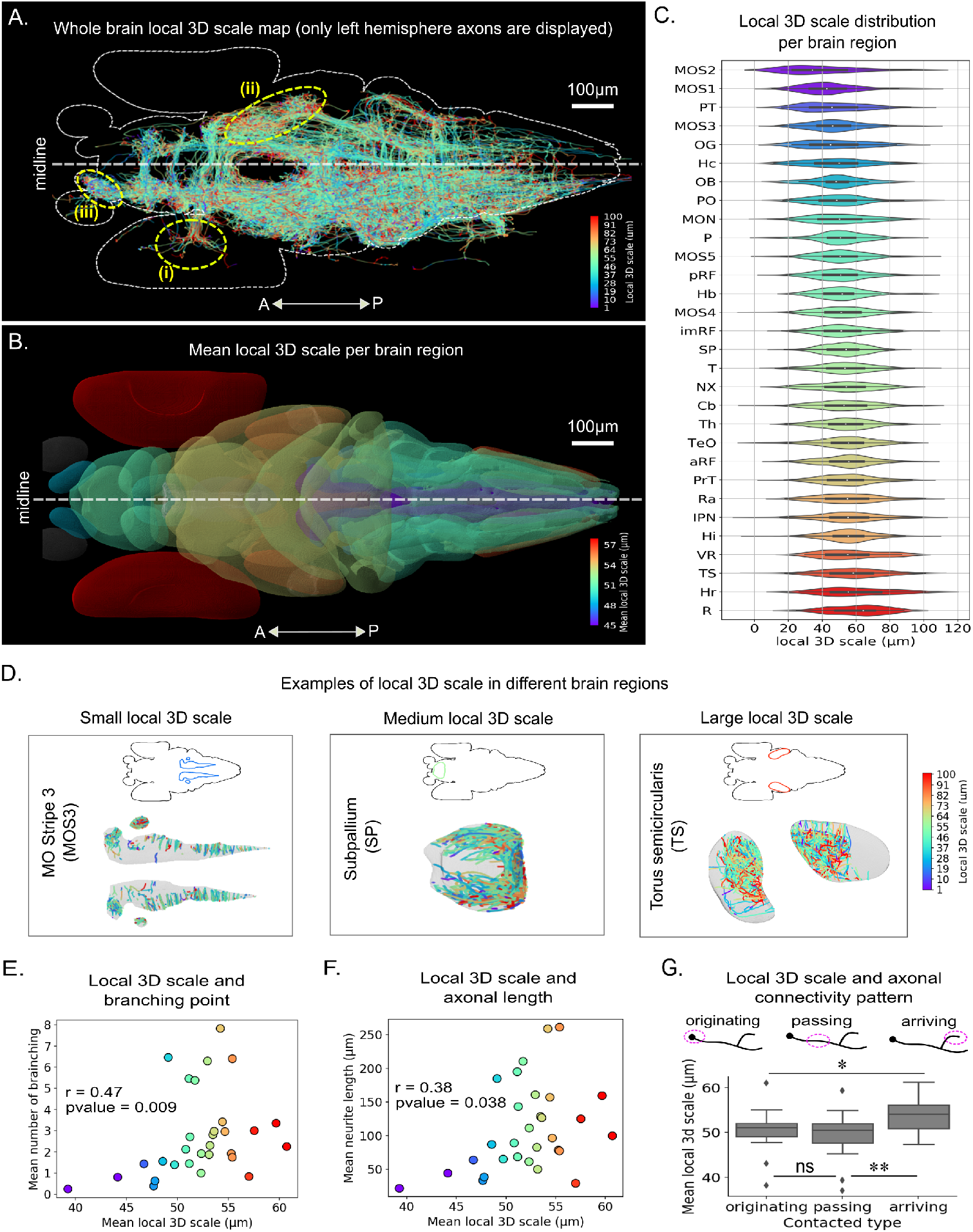
Local 3D scale mapping across the whole larval zebrafish brain. Traces analyzed correspond to those presented in Kunst et al. (2019). (A) Local 3D scale analysis of all axonal traces originating from neurons in the left hemisphere; the maximum scale is set to 100 *μ*m based on the width of the largest brain region. (B) Mean local 3D scales by brain regions. Regions with fewer than 15 traces are excluded (in gray). Values were clipped from the 5th to 95th percentiles for clearer display. (C) Distribution of the local 3D scale values in each brain region (see Table S1 for abbreviation definitions). (D) Examples of local 3D scale mapping for axonal traces in three brain regions respectively presenting low, medium and high mean 3D scale: the Medulla Oblongata Strip 3 (left), the Subpallium (middle), and Torus Semicircularis (right). (E, F) Correlation between the mean local 3D scales and average number of branching points (E) and trace length (F) in different brain regions. Spearman correlation was used. (G) Mean local 3D scale of axons originating from, passing through, and arriving to each brain region. Wilcoxon tests with Holm-Sidak correction for multiple comparison was used, * p≤0.05, ** p≤0.01, *** p≤0.005, **** p≤0.001. Color scale in C, E, F corresponds to that used in B.

To quantify these local differences, we computed the mean local 3D scale of inter region projections for each of the 36 brain regions defined by Kunst et al. This enabled us to establish an atlas of the local 3D scale of fish axons highlighting variations of their geometric complexity in different brain regions (Figure 3B and S7B). Similar atlases could be derived for all projection subsets originating from (Figure S8A), passing through (Figure S9A) or terminating in a region (Figure S10A), again showing inter-regional variations. The distributions of local 3D scale within each region nevertheless showed a high variance (Figure 3C, S8B, S9B and S10B), reflecting the diversity of neuronal types and variations in the local structure of their axons as they exit or enter a brain region. This variability is illustrated in Figure 3D showing three regions, the Medulla Oblongata Strip 3 (MOS3), Subpallium (SP) and Torus semicircularis (TS), respectively presenting low (~43 *μ*m), medium (~51 *μ*m) and high (~58 *μ*m) average local 3D scales. Different patterns exhibited by axons originating from, passing through and arriving in these regions might contribute to this variability (Figure S11).

Different distributions of local 3D scale could be observed in the fore-, mid- and hindbrain regions, with higher average value in mid- compared to fore- and hindbrain regions (Figure S7C). The difference is clearer for axons terminating in a region (Figure S10C), but not for axons with soma in these regions and en passant axons (Figures S8C, S9C). This could relate to the observation that the connectivity strength in the mid- and hindbrain are stronger than that in forebrain (Kunst et al., 2019).

Comparing the mean local 3D scale with the average number of branching points and average length of the traces within individual regions showed that these parameters were correlated, yet with a high variance (Figure 3E and 3F). However, when separately analyzing axons connecting to, through or from a region, only those originating from the region analyzed showed such correlations (Figures S8D-E), which were not apparent when observing passing or arriving axons (Figures S9D-E and S10D-E). Yet, comparing the local 3D scale of the three types of axons showed that inter-regional axons terminating in a region had on average higher local 3D scales than those originating from or passing through a region (Figure 3G). This observation is consistent with the idea that the proximal axonal path and terminal arbor differ in their complexity, as related to their developmental assembly and function.

Together, these results show that the local 3D scale computed with *nAdder* is informative at the whole brain level, adding a new level of description for analyzing connectomic datasets.

#### Characterization of axon behavior at defined anatomical locations and along specific tracts

A key feature of our metric is its local nature, i.e. its ability to inform on the geometry of single neurites not only globally, but at each point of their trajectory. This aspect is of interest to investigate the complexity of axonal trajectories not simply as function of their origin, but also at specific positions or spatial landmarks. For instance, commissural axons that cross the midline play a major role by interconnecting the two hemispheres and have been well studied, in particular for the specific way by which they are guided across the midline (Chédotal, 2014). Examining axonal arbors intersecting the midline in the whole larval fish dataset, we found that a majority (71%) crossed only once, making a one-way link between hemispheres, while a significant proportion crossed multiple times, with 10% traversing the midline more than 5 and up to 24 times (Figure 4A and S12A). Most individual branches (i.e. neuronal arbor segments between two branching points, or between a branching point and the cell body/terminal) crossed the midline just once, meaning that neurons crossing multiple times did not do so with one long meandering axons but with many distinct branches of their arbor (Figure S12B). Examples of such axonal arbors crossing the midline more than 10 times are shown in Figure 4A. Measuring the local 3D scale of crossing axon branches in a 20 *μ*m range centered on the midline (Figure 4B), we observed higher values for the ones traversing the midline several times compared to those only crossing once (Figure 4C, top). Axon branches crossing the midline multiple times were also much shorter than those crossing only once (Figure 4C, bottom). They localized mostly in the hindbrain in a ventral position (Figure 4D), while single crossings were frequent in the forebrain commissures. Finally, we found no link between the length and local 3D scale of crossing branches (Figure S12C). This is consistent with the existence of two populations of commissural axons in the vertebrate brain, the hindbrain being home to neurons following complex trajectories across the midline with many short and convoluted branches, and forebrain commissural neurons forming longer and straighter axons.

The local 3D scale metric can also serve to compare axon trajectories along entire tracts and characterize their average behavior. To illustrate this, we analyzed two distinct, well known cell types forming long-range axonal projections involved in sensory perception. We used *nAdder*, first studying the axon trajectories of retinal ganglion cells (RGCs) from the fish atlas, each originating in the retina and terminating in the tectum (Figure 4E). We observed that the local 3D scale of these axons presented clear heterogeneity across regions. To quantify this, we examined RGC axon trajectories in 5 portions of the retinotectal path: within the retina, at the retinal exit, at the midline, just prior to entering the tectum and within the tectum. We observed significant differences between these regions, with the mean local 3D scales at the midline presenting low values (~50 *μ*m) compared to the retina, the tectum, and their exit/entrance regions (~63 *μ*m). We then used *nAdder* to analyze a second population, the trajectories of Mitral cell axons originating in the olfactory bulb (OB) and projecting to multiple destinations in the pallium (P), subpallium (SP) or habenula (Hb) (Miyasaka et al., 2014). Here again we found variations in axonal complexity across regions (Figures 4F), with relatively low local 3D scales in the OB (~40 *μ*m) and Hb formation (~37 *μ*m) compared to higher values in the P (~50 *μ*m) and SP (~47 *μ*m). Overall, this indicates that local 3D scale is a relevant measure of geometric complexity at the single axon level, both to inform on specific points of their trajectory and to compare them within a neuronal class.

**Figure 4:**
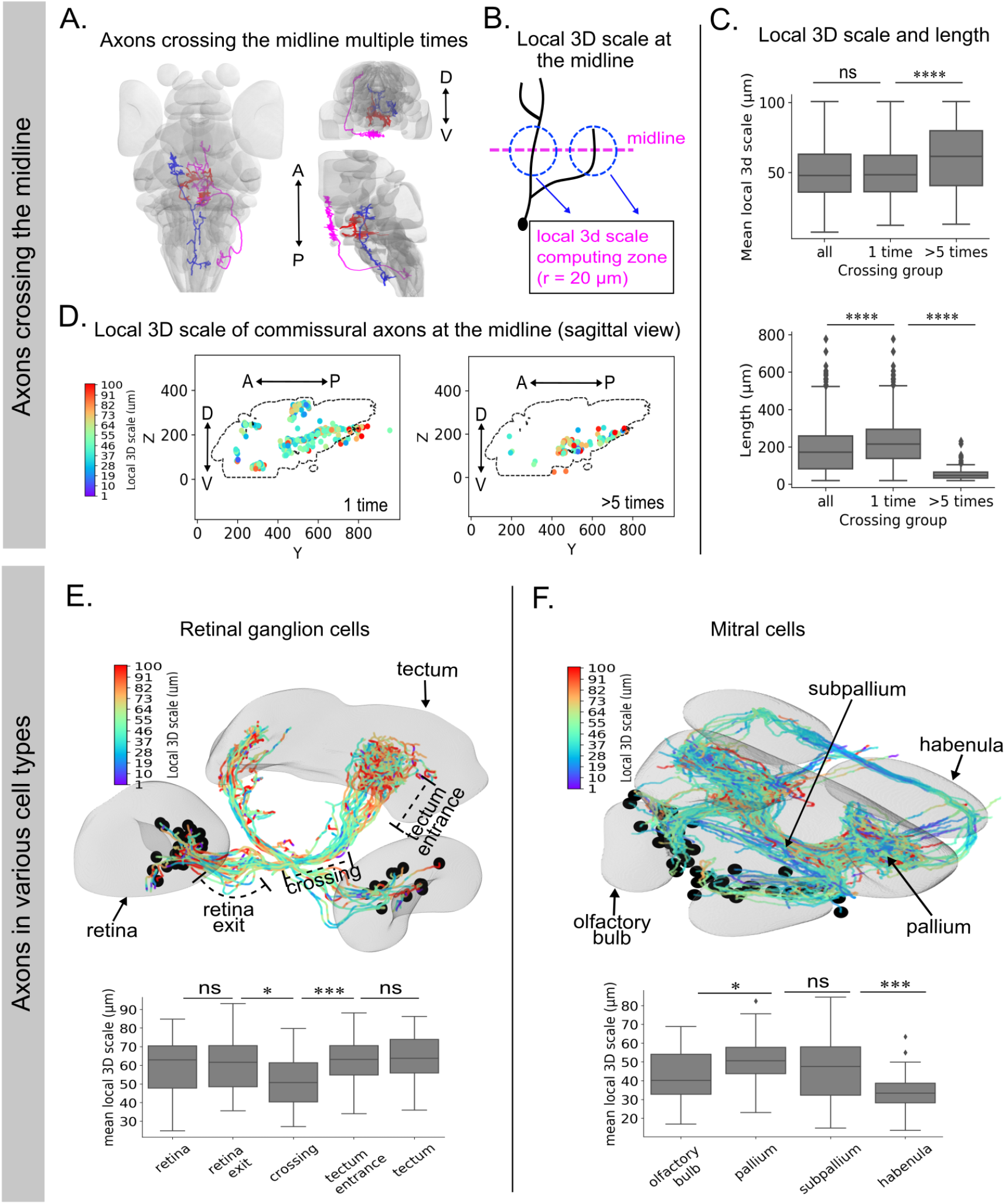
Local 3D scale analysis of specific locations and axonal tracts of the zebrafish brain. (A) Example of axonal arbors with many branches crossing the midline. (B) Diagram summarizing computation of the local 3D scale at the midline. (C) Local 3D scale (top) and length (bottom) of all axons crossing the midline vs. those crossing only once and more than five times. (D) View of the midline sagittal plane showing the local 3D scale of axons crossing only once and more than five times. The maximum scale is set to 100 *μ*m based on the width of the largest brain region analyzed. (E, F) Local 3D scale variation along axons of retina ganglion cells (E) and mitral cells (F). Wilcoxon tests with Holm-Sidak corrections for multiple comparison were used, * p≤0.05, ** p≤0.01, *** p≤0.005, **** p≤0.001.

##### Local 3D scale analysis of a brain region, the Torus Semicircularis

Finally, we investigated local 3D scale across different axonal populations innervating a given brain region. We performed this analysis in the Torus Semicircularis (TS) which presents a high average local 3D scale, with many complex axonal traces (Figure 3D right panel and S11 last row). Most of these traces corresponded to axons with terminations in the TS (70% vs. 5% axons emanating from TS and 25% axons passing through TS). Focusing on this population, we selected three groups of axons originating from the Tectum (TeO), Medial Octavolateral Nucleus (MON) and Medulla Oblongata Stripe 5 (MOS5), respectively, excluding other regions from which only a small number (< 10) of incoming axons had been traced (a detailed list of all regions with traces terminating in the TS is in Table S2). Axons coming from the TeO terminated in the ipsilateral TS (Figures 5A and 5B), in contrast to MON/MOS5 axons which connected the contralateral TS after crossing the midline (Figures 5D and 5E). Examining the distribution of local 3D scale inside the TS, we observed that tectal axons overall had a simpler structure, characterized by lower local 3D scale values than those coming from the MON/MOS5 (Figures 5C and 5F). In particular, complex axons with a local 3D scale above 150 *μ*m mostly originated from the MON/MOS5. This difference, revealed by *nAdder*, points to the existence of divergent developmental mechanisms of pathfinding and/or synaptogenesis for these two populations of axons within a shared target area.

**Figure 5:**
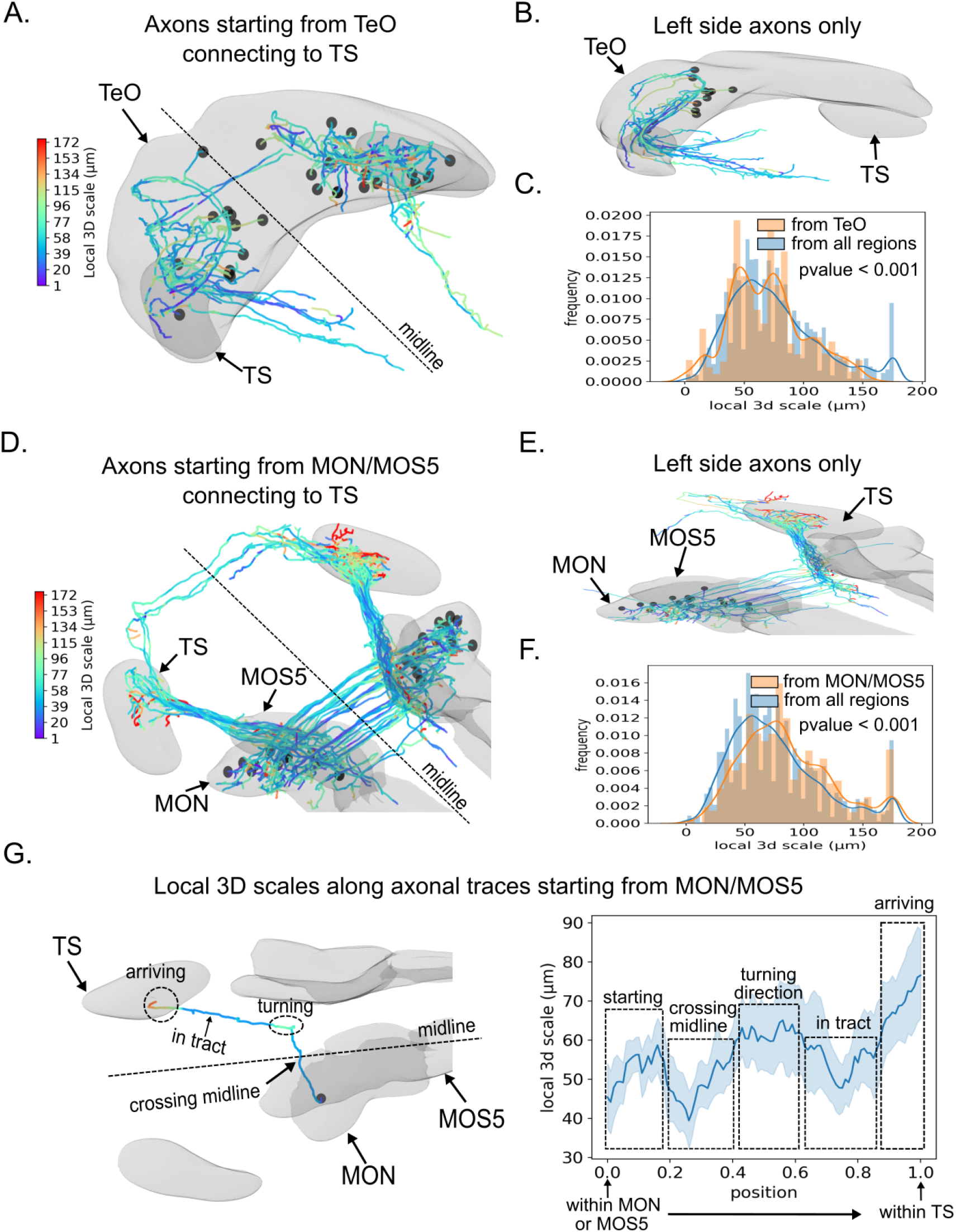
Local 3D scale of different axonal populations terminating in the zebrafish Torus Semicircularis (TS). (A, D) *nAdder* analysis of TS-terminating axons originating from the Tectum (TeO) (A) and Medial Octavolateral Nucleus (MON) or Medulla Oblongata Strip 5 (MOS5) (D). (B, E) Views restricted to axons originating from the left hemisphere. (C, F) Distribution of local 3D scale for the set of axons arriving from the TeO (C) and MON/MOS5 (F) compared to all axons arriving in the TS. Student’s t-test was used. (G) Local 3D scale variations along axons coming from MON/MOS5. Left, example view of one neuron indicating different segments of interest. Right, quantification (n = 32 axons). The maximum scale is set to 175 *μ*m based on the mean length of the longest branches. Only the longest branches are included in the computation.

We then focused on axons projecting from the MON/MOS5 to the TS. We computed the local 3D scale of these axons at all coordinates of their trajectory from the MON/MOS5 to the TS, normalized with respect to total length (Figure 5G). We found that these axons’ local 3D scale was highly correlated and that its average value varied widely along their course from the MON/MOS5 to the TS: low at the start, it increased inside the MON/MOS5 before dropping when the axons crossed the midline. It then increased again as they made a sharp anterior turn, dropped again within the straight tract heading to the TS, before rising to maximum values inside the TS. This result provides a striking example of stereotyped geometrical behavior among individual axons linking distant brain areas, likely originating from a same type of neurons, that our metric is able to pick up and report.

Overall, our analysis of the larval zebrafish brain atlas demonstrates that *nAdder* can be efficiently applied to connectomic datasets to probe structural heterogeneity and identify patterns among individual axons and population of neurons.

## 3 Materials and Methods

### 3.1 Intrinsic dimension decomposition of neurite branches based on scale-space theory

#### Intrinsic dimension decomposition

Let’s consider a 3D parametric curve *γ*(*u*) = (*x*(*u*)*, y*(*u*)*, z*(*u*)) for *u* ∈ [0, 1]; the *intrinsic dimensionality* of that curve can be defined as the smallest dimension in which it can be expressed without significant loss of information. An *intrinsic dimension decomposition* is a decomposition of a 3D curve into consecutive fragments of different intrinsic dimensionality, i.e. lying intrinsically on a 1D line, 2D plane or in 3D. Note that such decomposition needs not be unique. For example, not taking scales into account, two consecutive lines followed by one plane can in theory be decomposed into one 3D fragment, one 1D and one 3D fragments, two consecutive 1D and one 2D fragments, or two consecutive 2D fragments (Figure S1). The last two decomposition schemes are both meaningful and their combination represents a hierarchical decomposition of *γ* where linear fragments are usually parts of a larger planar fragment.

Here, we compute the decomposition of a curve intrinsically by looking at the curvature and torsion. Let us define the curvature *κ* of *γ* by *κ* = ||*γ′ × γ″||/||γ*′||^3^, corresponding to the inverse of the radius of the best approximation of the curve by a circle locally, the osculating circle. The torsion *τ* of *γ* is *τ* = ((*γ*′ × *γ*″) · *γ*‴)/||*γ*′ × *γ*″||^2^ and corresponds to the rate of change of the plane that includes the osculating circle. We then determine that if *κ* is identically equal to zero on a fragment then that fragment is a 1D line; similarly *τ* being identically equal to zero defines a 2D arc curve. Thus we define a linear indicator *L* of *γ*:

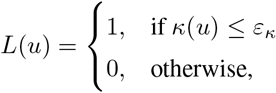

and a planar indicator *H* of *γ*:

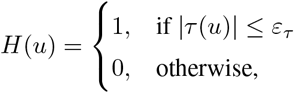

where *ɛ_κ_* and *ɛ_τ_* are the tolerances for computed numerical errors. We note that only using the *H* indicator is not sufficient to estimate the 2D plane. For example, a 2D plane fragment composed of a 1D line followed by a 2D arc cannot be entirely identified as 2D since the torsion is not defined as the curvature tends to 0. We therefore consider the planar-linear indicator *T* = *L* ∪ *H* instead of *H* for characterizing the curve according to such hierarchical order.

#### Scale space

In practice, the resulting dimension decomposition is closely tied to a ‘scale’ at which the curve is studied, i.e. up-close all diferentiable curves are well approximated by their tangent and thus would be linear. That property have been formalised through *Scale-spaces*, which have been described for curves in 2D (Mackworth & Mokhtarian, 1988) or 3D (Mokhtarian, 1988) in particular, to give a robust meaning to that intuition. A scale-space is typically defines as a set of curves *γ_s_* such that *γ_0_* = *γ* and increasing *s* leads to increasingly simplified curves. The scaled curve can be calculated by convolving *γ* with a Gaussian kernel of standard deviation *s* (Witkin, 1983), or by using mean curvature flow (Gage & Hamilton, 1986). In 2D, it has been shown for example that a closed curve under a mean curvature motion scale space will roundup with increasing s and eventually disappear into a point (Grayson, 1987).

However, the selection of a scale of interest representing a meaningful description of a curve is not straightforward. A scale defined as the standard deviation of the Gaussian kernel as above for example will be influenced by the kernel length and the curve sampling rate, and would be unintuitive to set. To make the scales become a more relevant physical/biological measurement, we employ as scale parameter the radius of curvature *r_κ_*, the inverse of the curvature *κ*, in *μ*m. For a given curve, the level of detail can be characterized by the maximal *r_κ_* along that curve, with scales keeping small *r_κ_* represents high level of detail looking at small objects such as small bumps and tortuosity and scales smoothing the curve out and keeing only larger *r_κ_* represents lower level of detail, and larger object and features such as plateaus or turns. Thus, to ease the interpretation and usage of scale spaces, we associate to a scale in micron 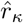 a set of standard deviation *s*, by determining, for a curve, for each point *u* whose 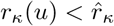 the standard deviation *s* such that 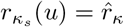. This therefore associates, for each curve, an anatomically relevant scale of interest in micron 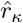 and the list *S* of standard deviations that lead to that curve having no points with curvature radius smaller than 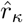. We will then use *S* to estimate the dimension decomposition of *γ* at scale 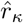 in two steps, first by computing our indicators for all *s* ∈ *S*, then by selecting, across *S*, the best decomposition.

First we compute *L_s_* and *T_s_* by calculating the curvatures *κ_s_* and the torsions *τ_s_*. We then exclude fragments smaller than a length threshold *ɛ_ω_* at every *s* to eliminate small irrelevant fragments. From *L_s_* and *T_s_*, the linear fragments denoted as 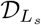 and planar-linear fragments denoted as 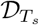 are deduced across all *s* ɛ *S*. An example of this sequence of dimension decompositions is illustrated in Figure S4 where 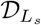 and 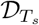 are superimposed. The curve is completely 3D at low *s* value, several linear and planar fragments then appear at higher *s*, progressively fuse to form larger linear and planar fragments and eventually converge to one unique line at very high value of *s*.

In the second step, we estimate the most durable combination of fragments from the linear segments 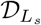 and linear-planar segments 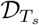 calculated in the first step. We compute the number of fragments at every *s* ∈ *S*, to measure how long each combination of fragments exists. We then select the combination of fragments remaining for the longest subinterval of *S*. Knowing that each fragment in the combination can have different lengths among that subinterval, we thus select the longest candidate. In cases where fragments overlap, we split the overlap in half. Of note, we repeat this step twice, first to estimate the best combination of linear-planar segments from 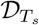 input and mark the corresponding subinterval, then to estimate the best combination of linear fragments from 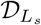 input within that subinterval. As seen in Figure S4, the curve is decomposed at every *s* in consecutive planar/nonplanar fragments, then the linear fragments are identified within each planar fragment. The final result is the hierarchical dimension decomposition of the curve *γ* at a given scale of interest 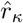.

### 3.2 Evaluation of dimension decomposition on simulated curve

The simulated curve consists of consecutive 1D lines, 2D planes and 3D regions. The simulation of 1D lines is done by simply sampling a sequence of points on an arbitrary axis (e.g. *x* axis), then rotating it in a random orientation in 3D. The simulation of random yet regular 3D fragments embedded in a 2D plane is more challenging. We tackle this issue by using the active Brownian motion model (Volpe et al., 2014). Active Brownian motion of particles is an extended version of the standard Brownian motion (Uhlenbeck & Ornstein, 1930) by adding two coefficients the translational speed to control directed motion and the rotational speed to control the orientation of the particles. At each time point, we generate the new coordinates (*x, y,* 1) by the following formulas:

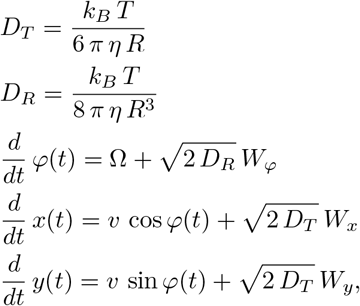

where *D_T_* and *D_R_* are the translational and rotational coefficients, *k_B_* the Boltzmann constant, *T* the temperature, *η* the fluid viscosity, *R* particle radius, *φ* rotation angle, Ω angular velocity and *W_φ_, W_x_, W_y_* independent white noise. After generating the simulated intrinsic 2D fragment, a random rotation is applied. For simulating a 3D fragment, we extended the Active Brownian motion model in 2D (Volpe et al., 2014) to 3D as follows:

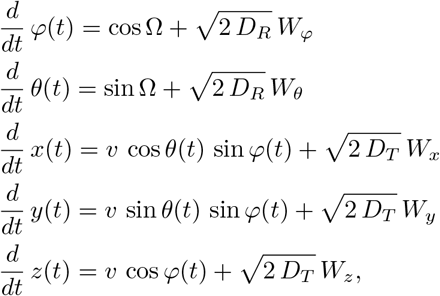

where (*φ, θ*) are spherical angles. We simulate the curve *γ* with a sequence of *n_ω_* consecutive fragments of varying random intrinsic dimensions. The curve *γ* is then resampled equally with *n_γ_* points and white noise 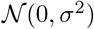 is added.

We set *n_ω_ ≤* 5, *n_γ_* = 1000 points and vary *σ* between 1 and 30 *μm*. The upper value of *σ* = 30 *μm* is high enough to corrupt local details of a simulated fragment with about 100 *μm* of length in our experiment.

The metric used to measure the accuracy of the intrinsic dimension decomposition, defined to be between 0.0 and 1.0 and measures an average accuracy across fragments, is given by 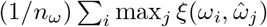, where 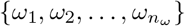 are the simulated intrinsic fragments, 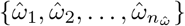 the estimated intrinsic fragments and *ξ*(.,.) the *F*_1_ score (Rijsbergen, 1979) calculated by:

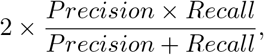

where

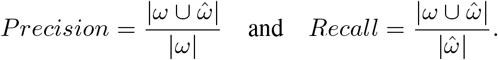

In order to evaluate our intrinsic dimensions decomposition algorithm, we look at the curve *γ* at various scales *r_κ_* = 1, 2*,…,* 100 *μm*, then perform the dimension decomposition for every *r_κ_*. Setting *r_κ_* = 100 *μm* is large enough to get a smoothed curve whose simulated intrinsic fragments are no longer preserved. In addition, we also removed estimated fragments whose lengths were smaller than *ɛ_ω_*. Here we set *ɛ_ω_* = 5% of the curve length since it does not surpass the minimal length of simulated intrinsic fragments. While automatic determination of an optimal scale is not studied in this paper, with the *nAdder* algorithm being in part meant to sidestep this complex issue altogether by computing a single metric across scales, we have to chose a scale for the evaluation. We either take the one corresponding to the largest accuracy (optimal scale, Figure S2) or one at a fixed scale (Figure S3), *r_k_* = 20 *μm*, small enough to avoid deforming the simulated curve. Our method is then compared with a baseline method (Yang et al., 2016; Ma et al., 2017) that first iteratively assigns each point on the curve as linear/nonlinear by a collinearity criterion, then characterize nonlinear points as planar/nonplanar by a coplanarity criterion. For both methods, the curve was first denoised based on Czesla et al. (2018).

### 3.3 Local 3D scale computation

Assuming a scale interval of interest, the local 3D scale of a point on a curve is computed by iteratively checking at each scale the intrinsic dimension of that point and taking the scale at which the local scale is not 3D anymore. In some cases, the dimension transformation of a point is not smooth, since the point at a 2D/1D scale can convert back to 3D. We therefore select the first scale of the longest 2D/1D subinterval as local 3D scale. In case no subintervals are found, it means that the point remains 3D across the whole interval and its local 3D scale is set as the highest scale.

### 3.4 Local 3D scale analysis of neuronal traces

We consider a neuronal trace as a 3D tree and first resample it with a spacing distance between two points equal to 1 *μ*m. B-spline interpolation (Boor, 1978) of order 2 is used for the resampling. We then decompose the neuronal arbor into potentially overlapping curves, since the intrinsic dimension decomposition and local 3D scale are computed on individual neuronal branches. Several algorithms are available for tree decomposition. We explored three, functioning either by longest branch (first taking the longest branch, then repeating this process for all subtrees extracted along the longest branch), by node (taking each curve between two branching nodes) or by leaf (taking, for each leaf, the curve joining the leaf to the root). Local scales were computed for all three considered decompositions and the ‘leaf’ decomposition was found to give the most robust and meaningful results as shown in Figure S5. That method gives partially overlapping curves; thus, we compute the average of the values when needed; we verified that the standard deviations were small.

Next, the local 3D scale is computed for each extracted sub-branch within a scale interval of interest. The scale is defined as the radius of curvature and can be selected based on the size of the branches or that of the region studied. For example, in the case of Xeonopus Laevis tadpole neurons (Figure 2), we selected scales from 1 to 60 *μ*m which was sufficient to study neuronal branches of about 180 *μ*m in length (i.e. a 60 *μ*m radius of curvature corresponds to a semicircle with length equal to 60*π ≈* 180 *μ*m). In the case of the zebrafish brain dataset (Figure 3), we selected scales from 1 to 100 *μ*m based on the length of the largest region, which was about 300 *μ*m. When studying axons arriving to the Torus Semicircularis (Figure 5), we set the maximum scale to 175 *μ*m as the neuronal branches are on average 525 *μ*m long. Moreover, we excluded the sub-branches whose lengths are smaller than a threshold set to 5 *μ*m, which is half the size of the smallest brain region. Branches shorter than 5 *μ*m contribute very little to the local 3D scale result. This is also useful to reduce artifact caused by manual tracing of neurons.

### 3.5 Toolset for neuronal traces manipulation

We provide an open source Python toolset for seamless handling of 3D neuronal traces data. It supports reading from standard formats (swc, csv, etc), visualizing, resampling, denoising, extracting sub-branches, calculating basic features (branches, lengths, orientations, curvatures, torsions, Strahler order (Horton, 1945), etc), computing the proposed intrinsic dimension decompositions and the local 3D scales. The toolset is part of the open source Python library GeNePy3D (Phan & Chessel, 2020) available at https://genepy3d.gitlab.io/ which gives full access to manipulation and interaction of various kinds of geometrical 3D objects (trees, curves, point cloud, surfaces).

### 3.6 Analyzed neuronal traces datasets

The traces used in our studies are all open and available online for downloading. They consist of neuronal reconstructions from NeuroMorpho.Org and the Max Planck Zebrafish Brain Atlas listed in the table below:

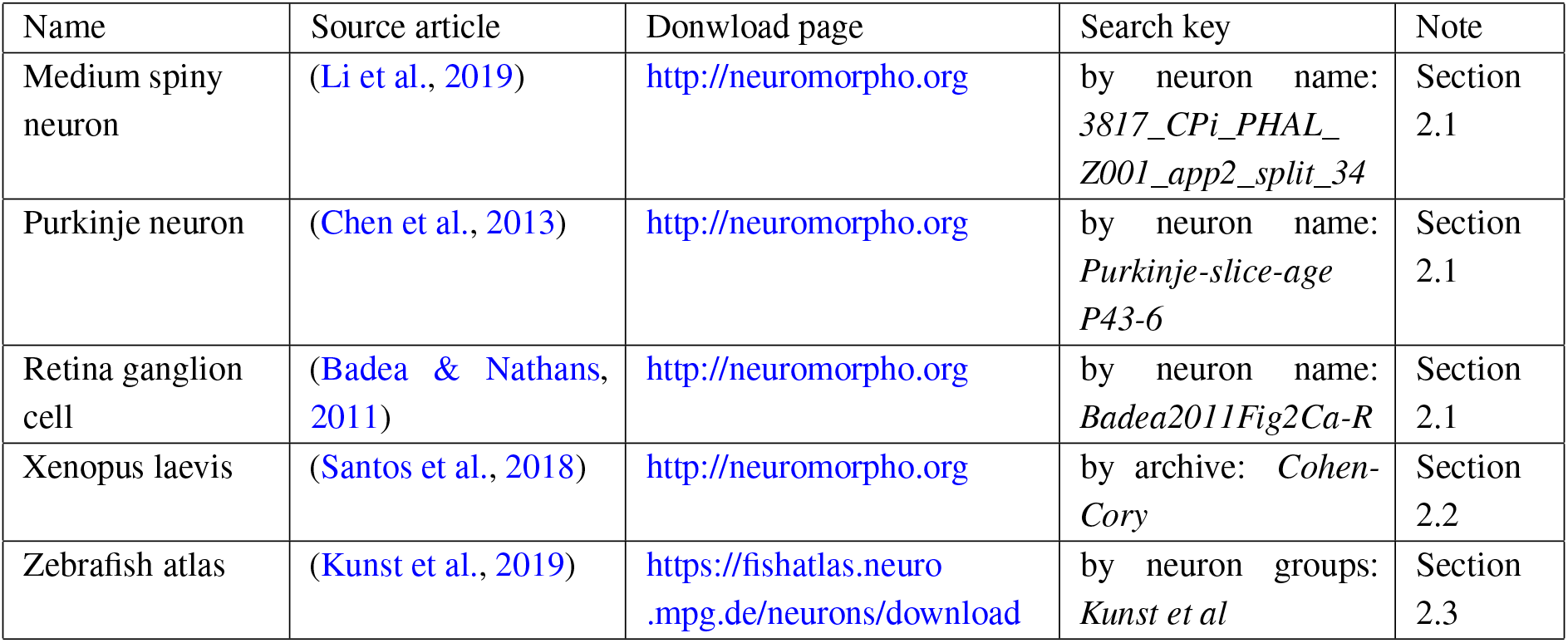

| Name | Source article | Donwload page | Search key | Note |
| --- | --- | --- | --- | --- |
| Medium spiny neuron | (Li et al., 2019) | http://neuromorpho.org | by neuron name: *3817_CPi_PHAL_Z001_app2_split_34* | Section 2.1 |
| Purkinje neuron | (Chen et al., 2013) | http://neuromorpho.org | by neuron name: *Purkinje-slice-age P43-6* | Section 2.1 |
| Retina ganglion cell | (Badea & Nathans, 2011) | http://neuromorpho.org | by neuron name: *Badea2011Fig2Ca-R* | Section 2.1 |
| Xenopus laevis | (Santos et al., 2018) | http://neuromorpho.org | by archive: *Cohen-Cory* | Section 2.2 |
| Zebrafish atlas | (Kunst et al., 2019) | https://fishatlas.neuro.mpg.de/neurons/download | by neuron groups: *Kunst et al* | Section 2.3 |

## 4 Discussion

We presented a novel method for the multiscale estimation of intrinsic dimensions along an open 3D curve and used it to compute a local 3D scale, which we propose as a new local metric to characterize the geometrical complexity of neuronal arbors. We demonstrated that our method, dubbed *nAdder*, is accurate and robust, and applied it to published trace data, showing its relevance and usefulness to compare neurite trajectories from the level of single neurons to the whole brain.

Our new approach provides a solution to the problem of measuring the geometrical complexity of 3D curves not just globally but also locally, i.e. at successive points of their trajectory. It enables to compute the dimensionality of such curves at any position, for a given scale of analysis. By scanning the scale space, one can then determine the value at which the curve requires all three dimensions of space for its description, a sampling-independent local metric which we term the local 3D scale. Mathematically, scale spaces of curves are trickier to implement in 3D than in 2D and have been much less studied. In 2D for example, scale spaces based on curvature motion are known to have advantageous properties compared to Gaussian ones: they arise naturally from principled axiomatic approaches, are well described mathematically and can be computed through well characterized numerical solutions of partial differential equations (Cao, 2003). They are readily extended to 3D surfaces by using principal curvatures (Brakke, 2015), but no simple equivalent is known for 3D curves. One theoretical issue is that using only the curvature to determine the motion of 3D curves would not affect the torsion, and for example a set of increasingly tighter helices would have increasingly higher curvature but identical torsion. Here, we provided and thoroughly evaluated a practical solution to this problem by associating a spatial scale to an ensemble of Gaussian kernels. In future work, additional theoretical studies of 3D scale spaces of curves could potentially lead to simpler and more robust algorithms with solid mathematical foundations.

The *nAdder* approach fills a gap in the methodologies available to analyze neuronal traces, which until now were lacking a straightforward way to measure their geometrical complexity and its variations, both within and between traces. It is complementary to techniques classically used to study the topology of dendritic and axonal arbors (number and position of branching points, for example), providing a geometrical dimension to the analysis that is a both simpler and more robust than the alternative direct computation of curvature and torsion. In practice, the local 3D scale of neurons (as measured with nAdder and observed in this study) ranges from 0 to 100-200 *μ*m. The lowest values are typically found in axonal tracts that follow straight trajectories, or in special cases such as the planar dendritic arbor of cerebellar Purkinje neurons shown in Figure 1C2. Higher, intermediate values are for instance observed at inflexion points along tracts where axons change course in a concerted manner. The highest values of local 3D scale typically characterize axonal arbor portions where their branches adopt a more exploratory behavior, in particular in their most distal segments where synapses form. A striking example is given in Figure 5G, showing the trajectory of an axon from the MON/MOS5 to the torus semicircularis, where changes in local 3D scale correlate with distinct successive patterns: a straight section through the midline, a sharp turn followed by another straight section before more complex patterns in the target nucleus. Importantly, nAdder not only enables to quantify how geometric complexity varies along a single axon trace, but also to compare co-registered axons of a same type and to characterize coordinated changes in their behavior (Figure 5G, right panel). It thus offers a way to identify key points and stereotyped patterns along traces. This will be of help to study and hypothesize on the developmental processes at the origin of these patterns, such as guidance by attractive or repulsive molecules (Lowery & Van Vactor, 2009), mechanical cues (Gangatharan et al., 2018), or pruning (Lichtman & Colman, 2000; Riccomagno & Kolodkin, 2015).

The high values of local 3D scale near-systematically observed in the most distal part of axonal arbors are also of strong interest, as these segments typically undergo significant activity-based remodeling accompanied by branch elimination during postnatal development (Lichtman & Colman, 2000), resulting in complex, convoluted trajectories in adults (Keller-Peck et al., 2001; Lu et al., 2009). In accordance with this, we observed a correlation between local 3D scale and axon branching at the whole brain level (Figure 3E). Further integrating topological and geometrical analysis and linking these two aspects with synapse position could then be a very beneficial, if challenging, extension of *nAdder*. To this aim, one would greatly benefit from methods enabling both faithful multiplexed axon tracing over long distances and mapping of the synaptic contacts that they establish. Progress in volume electron (Motta et al., 2019), optical (Abdeladim et al., 2019) or X-ray holographic microscopy (Kuan et al., 2020) will be key to achieve this in vertebrate models.

Beyond neuroscience, we also expect *nAdder* to find applications in analyzing microscopy data in other biological fields wherever 3D curves are obtained. For example, it could help characterizing the behavior of migrating cells during embryogenesis (Faure et al., 2016), or be used beside precise biophysical models to interpret traces from single-particle tracking experiments (Shen et al., 2017).

Importantly, an implementation of the proposed algorithms, along with all codes to reproduce the figures, is openly available, making it a potential immediate addition to computational neuroanatomy studies. Specifically, *nAdder* is part of GeNePy3D, a larger quantitative geometry Python package providing access to a range of methods for geometrical data management and analysis. More generally, it demonstrates the interest of geometrical mathematical theories such as spatial statistics, computational geometry or scale-space for providing some of the theoretical concepts and computational algorithms needed to transform advanced microscopy images into neurobiological understanding.

## Acknowledgments

This work was supported by Agence Nationale de la Recherche (contract ANR-11-EQPX-0029 Morphoscope2) as well as Fondation pour la Recherche Médicale (DBI20141231328) and Agence Nationale de la Recherche under contracts LabEx LIFESENSES (ANR-10-LABX-65) and IHU FOReSIGHT (ANR-18-IAHU-01).

