## Supplemental Figures for "nAdder: A scale-space approach for the 3D analysis of neuronal traces"

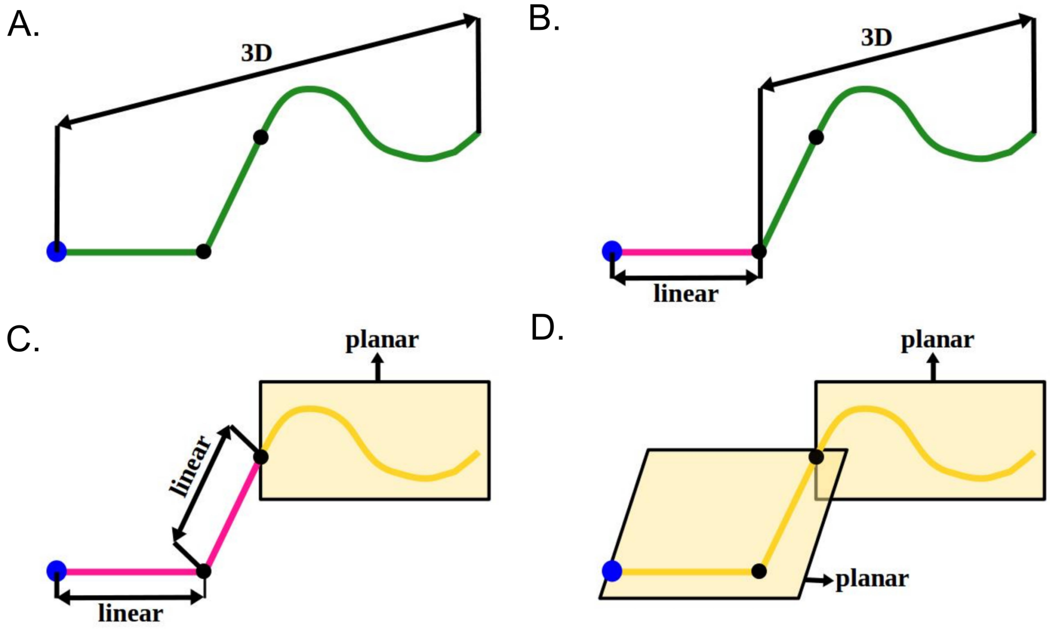


**Figure S1: Different schemes for the intrinsic decomposition of 3D traces.** (A) the trace is entirely 3D. (B) the trace is decomposed by 1D line followed by 3D fragment. (C) the trace is decomposed by suite of 1D lines and 2D plane. (D) The trace is decomposed by suite of 2D planes. The decompositions are hierarchial where a 1D line is lying on a 2D plane, and the 2D plane itself is lying within a 3D portion.

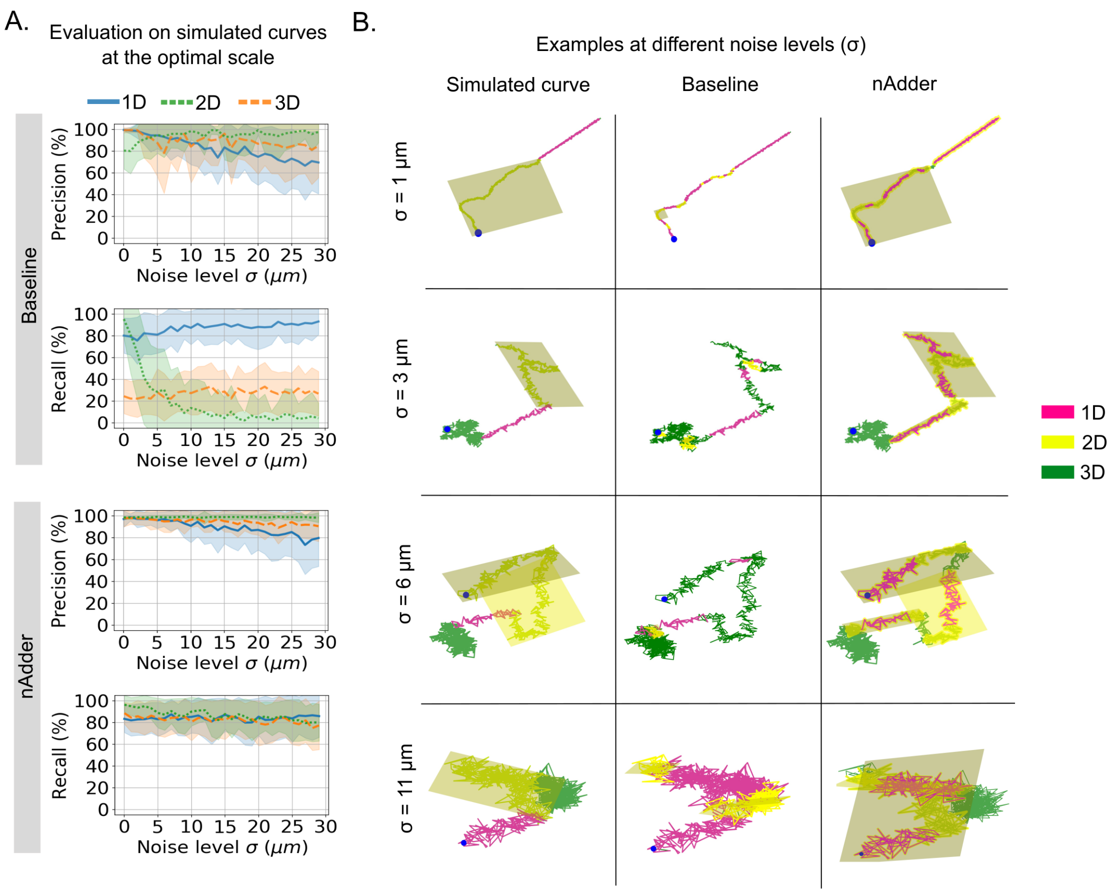


**Figure S2: Evaluation of intrinsic dimensional decomposition on simulated 3D traces.** (A) Precision and Recall of the proposed algorithm nAdder and the baseline (Yang *et al.*, 2016; Ma *et al.*, 2017) as a function of the noise level $\sigma$ varying between 1 and 30 $\mu$m. The algorithms are applied at various scales from 1 to 100 $\mu$m and the scale with the largest accuracy (optimal scale) is chosen. (B) Comparision of estimated intrinsic dimensions at four different noise levels. Compared with the baseline approach, the nAdder is more robust to noise and gives much higher accuracies in both Precision and Recall. Details of the algorithms and simulations are shown in Methods.

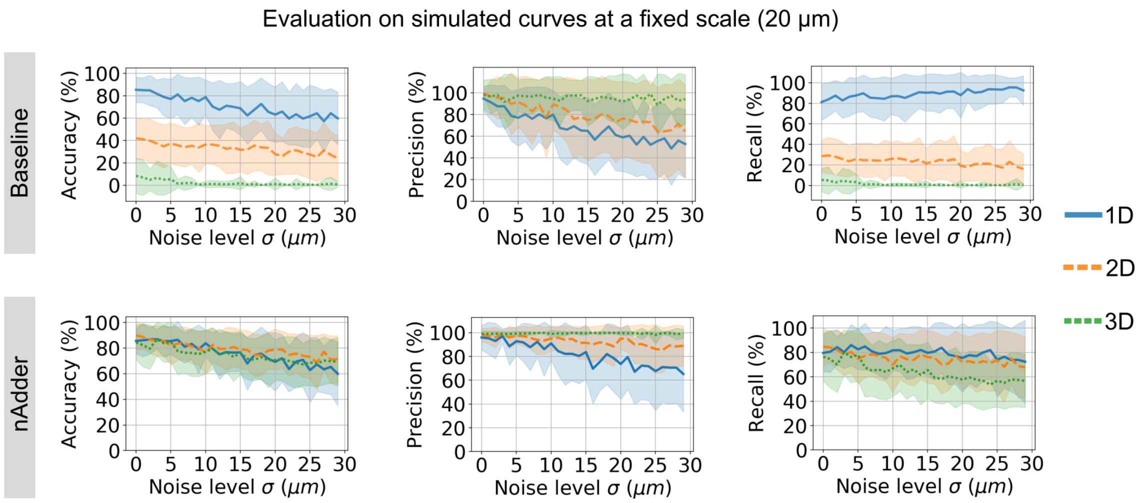


**Figure S3: Evaluation of the dimensionality decomposition algorithm at a fixed scale.** Accuracy, Precision and Recall of the nAdder and the baseline approach from Yang *et al.* (2016); Ma *et al.* (2017) as a function of the noise level $\sigma$ varying between 1 and 30 $\mu$m. The algorithms are applied at a fixed scale = 20 $\mu$m, which is small enough to avoid deforming the simulated curve. The accuracies of both algorithms are not as high as in the case of an optimal scale (Figure S1), but our approach still acheives ~85% of accuracy at $\sigma$ = 5 (medium noise) and ~80% of accuracy at $\sigma$ = 10 (high noise) for both 1D, 2D and 3D, compared to much lower accuracies for the baseline in 2D and 3D.

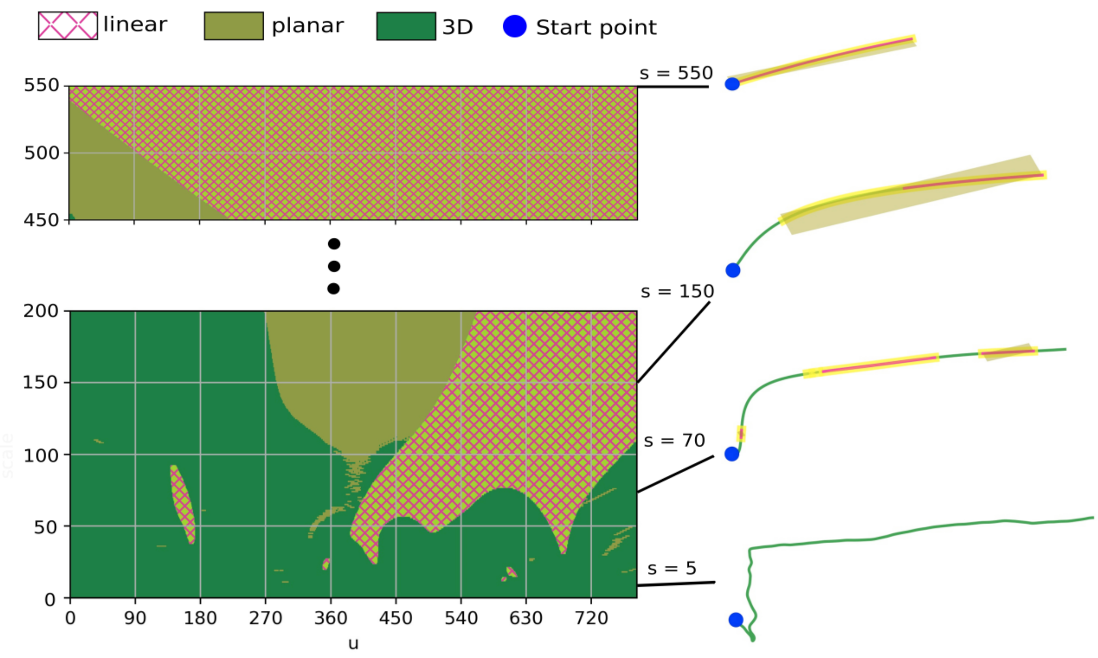


**Figure S4: Intrinsic dimensional decomposition of an axonal trace across multiple scales.** Positions on the 3D trace are indexed by $u$ (column of left panel) and the scaled trace is calculated by Gaussian convolution with various standard deviations (rows in the left panel). An example of trace seen at different scales and superimposed with its decomposition is shown on the right. The trace exhibits mostly 3D at small scales, then decomposes into a combination of 1D/2D/3D portions at higher scales and finally transforms into a 1D line at a very high scale.

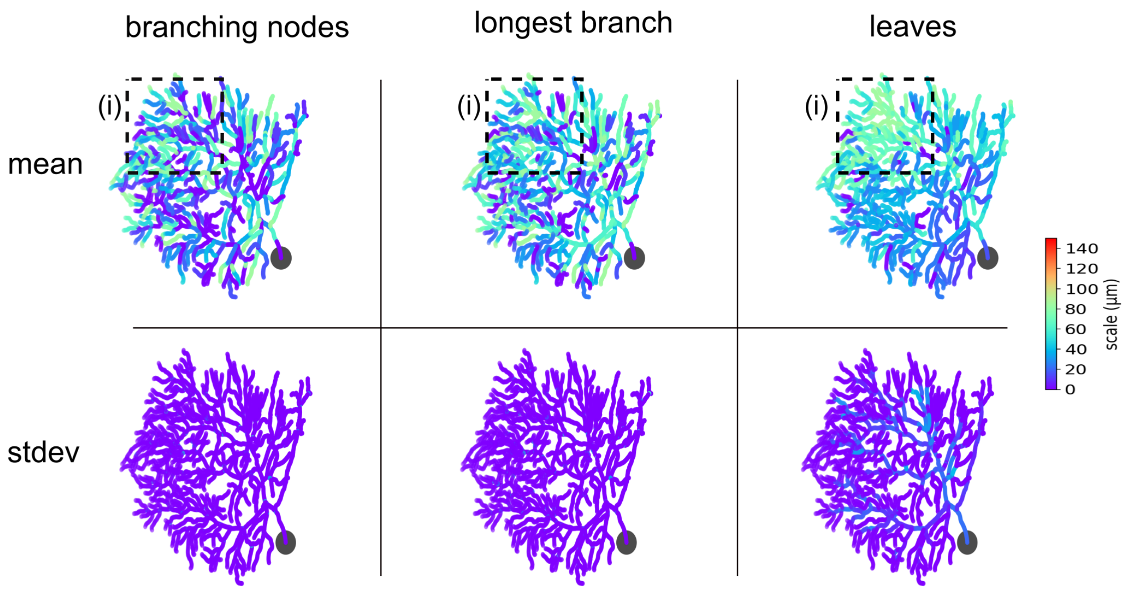


**Figure S5: Computation of local 3D scales in different decomposition modes on the cerebellar Purkinje neuron shown in Figure 1C2.** (Left) neurite portions located between any two branching nodes were extracted. (Middle) the longest branch originating from the cell body (root tree) was first extracted, and the process repeated for all subtrees extracted from that longest branch. (Right) branches connecting the cell body to each leaf were extracted. The mean and standard deviation of the local 3D scale was computed. The "leaves" mode (right) produces more stable local 3D scales with high and homogenous values in region (i) where the dendrites are sticked out of plane compared to the "branching nodes" (left) and "longest branch" modes (middle) (See Movie S2).

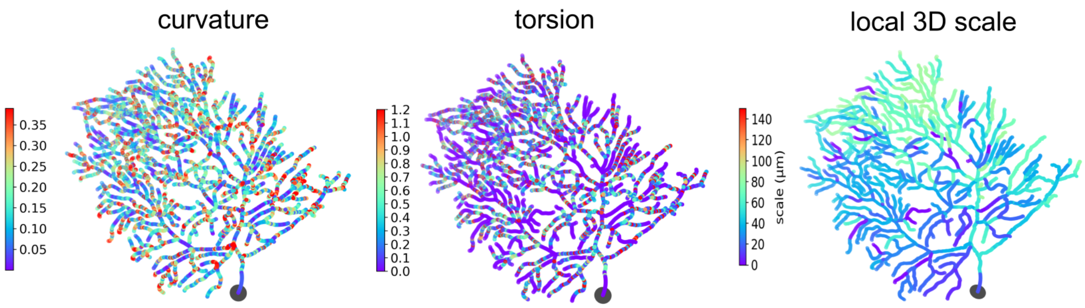


**Figure S6: Comparison between different local metrics of geometrical complexity.** Three parameters, curvature (left), torsion (middle) and local 3D scale (right) were mapped on the cerebellar Purkinje neuron shown in Figure 1C2. The local 3D scale gives smoother values than those of curvature and torsion since it was computed using a scale space approach, and better contrasts different regions of the Purkinje cell's dendritic arbor.

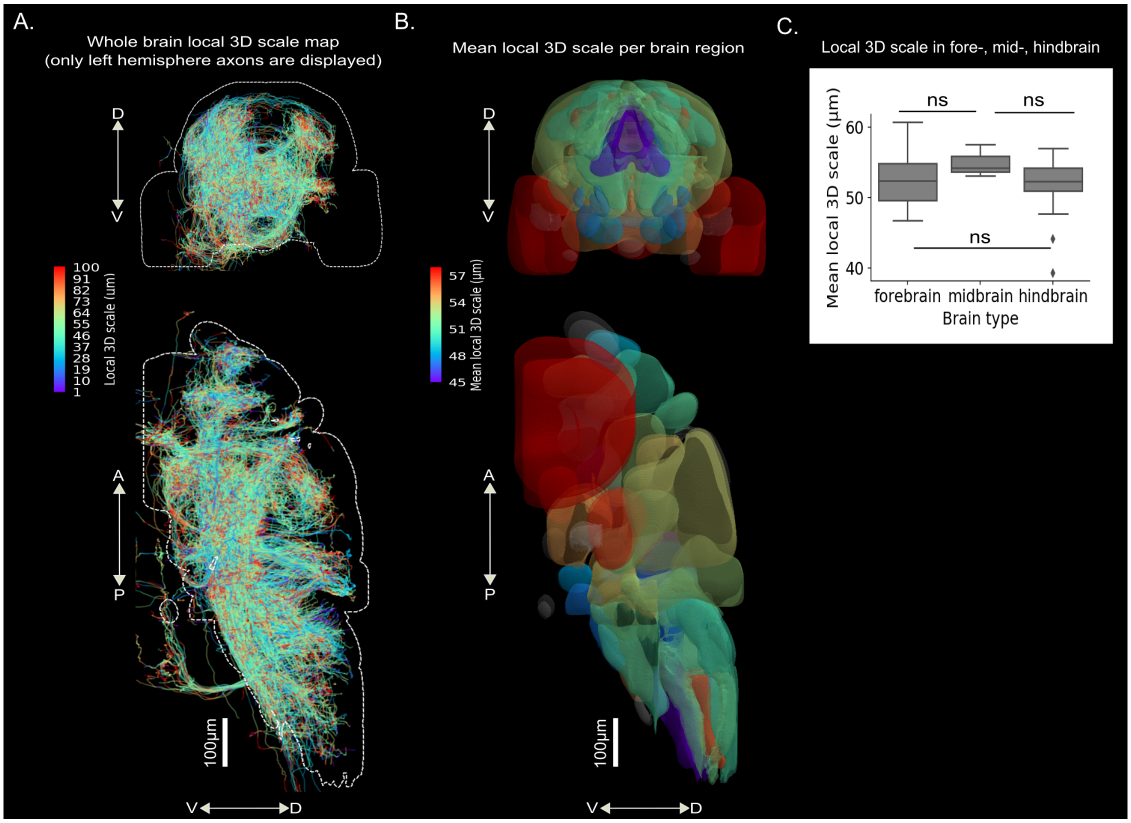


**Figure S7: Local 3D scales mapping across the whole larval zebrafish brain in front and side views.** The traces analyzed correspond to those presented in Kunst *et al.* (2019). (A) Local 3D scale analysis of all axonal traces originating from left hemisphere. (B) Mean local 3D scale by brain regions (values were clipped from $5^{th}$ to $95^{th}$ percentiles for clearer display). A transparency effect was applied to help visualizing inter-regional variations. (C) Mean local 3D scale in the fore- mid- and hindbrain. A Wilcoxon test with Holm-Sidak for multiple comparison was used, ns = not significant.

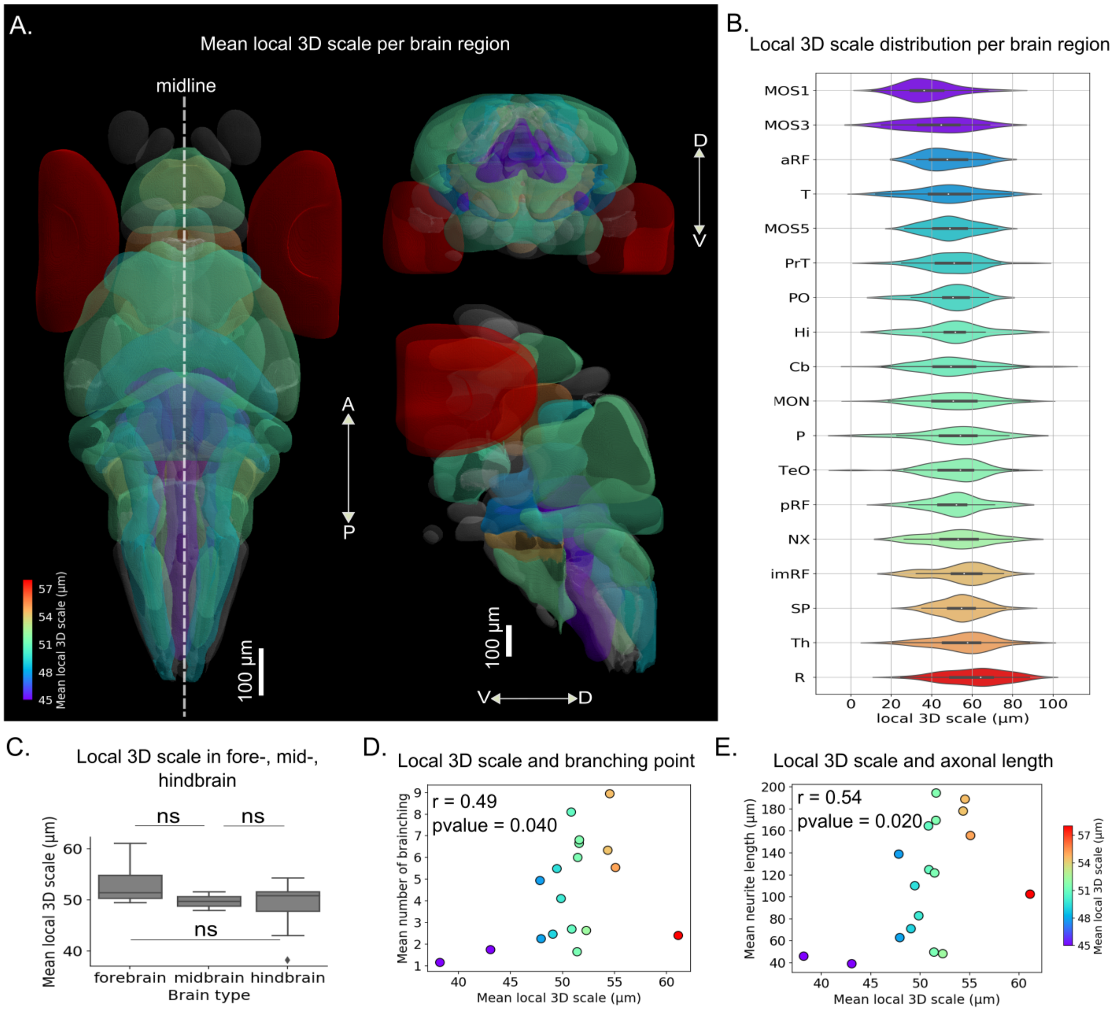


**Figure S8: Whole brain local 3D scale analysis of axons originating from different region of the larval zebrafish brain.** Traces analyzed correspond to those presented in Kunst *et al.* (2019). (A) Mean local 3D scale by brain region (values were clipped in the same range as in Figure S6B for comparison). (B) Distribution of the local 3D scale values in each brain region. (C) Mean local 3D scale in fore- mid- and hindbrain. A Wilcoxon test with Holm-Sidak for multiple comparison was used, ns = not signficant. (D, E) Correlation between the mean local 3D scales and average number of branching points (D) and trace length (E) in different brain regions. Spearman correlation was used.

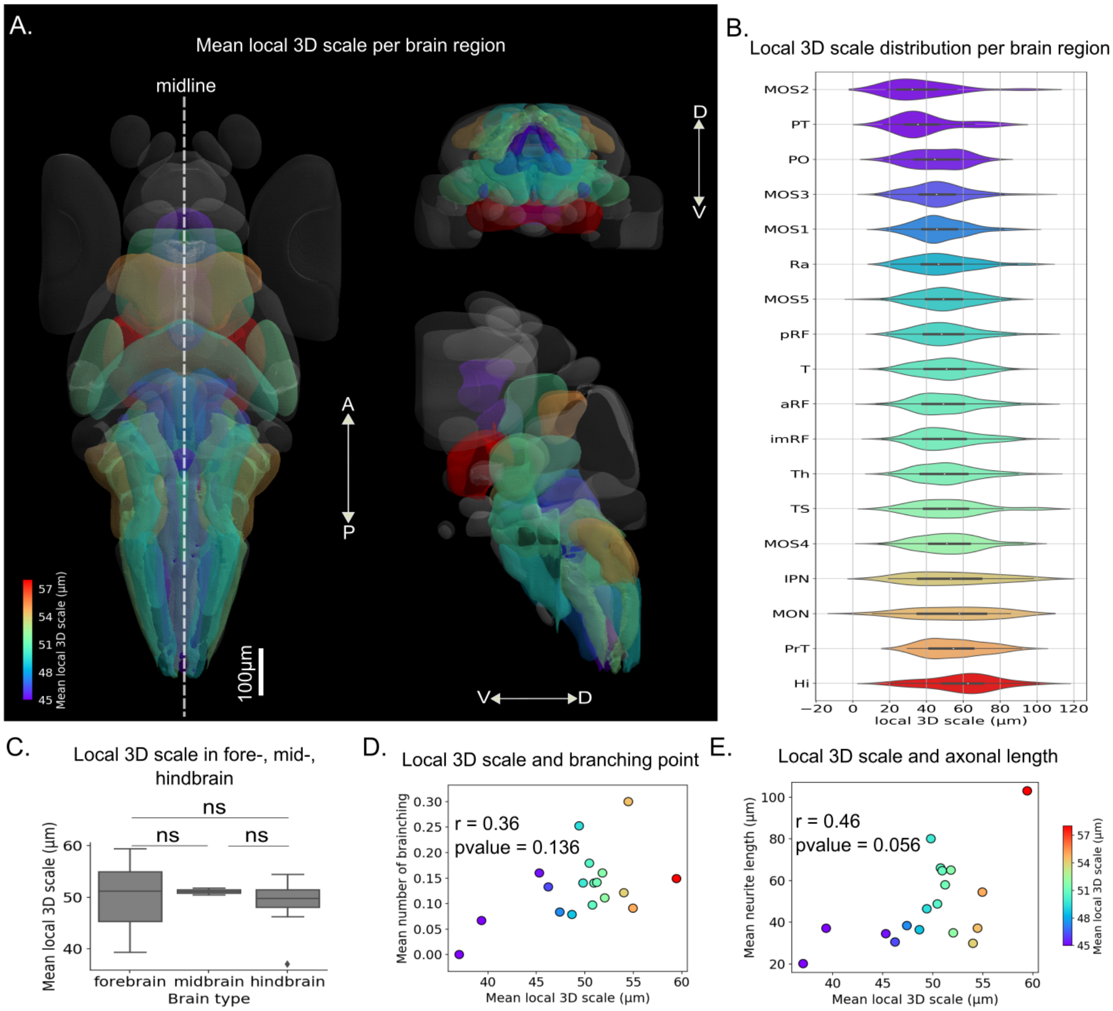


**Figure S9: Whole brain local 3D scale analysis of axons passing through different region of the larval zebrafish brain.** Traces analyzed correspond to those presented in Kunst *et al.* (2019). (A) Mean local 3D scale by brain region (values were clipped in the same range as in Figure S6B for comparison). (B) Distribution of the local 3D scale values in each brain region. (C) Mean local 3D scale in fore- mid- and hindbrain. A Wilcoxon test with Holm-Sidak for multiple comparison was used, ns = not signficant. (D, E) Correlation between the mean local 3D scales and average number of branching points (D) and trace length (E) in different brain regions. Spearman correlation was used.

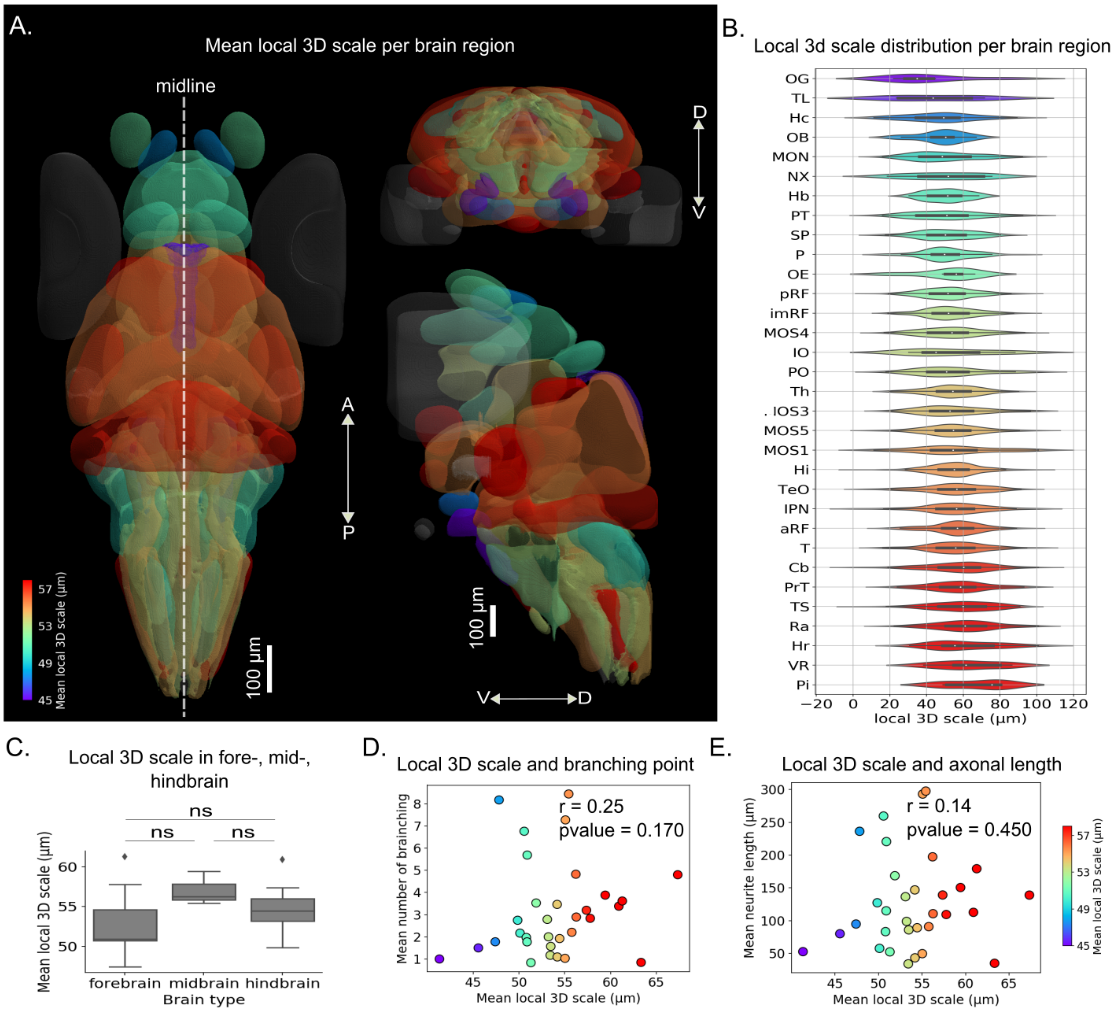


**Figure S10: Whole brain local 3D scale analysis of axons terminating in different region of the larval zebrafish brain.** Traces analyzed correspond to those presented in (Kunst *et al.*, 2019). (A) Mean local 3D scale by brain region (values were clipped in the same range as in Figure S6B for comparison). (B) Distribution of the local 3D scale values in each brain region. (C) Mean local 3D scale in fore- mid- and hindbrain. A Wilcoxon test with Holm-Sidak for multiple comparison was used, ns = not signficant. (D, E) Correlation between the mean local 3D scales and average number of branching points (D) and trace length (E) in different brain regions. Spearman correlation was used.

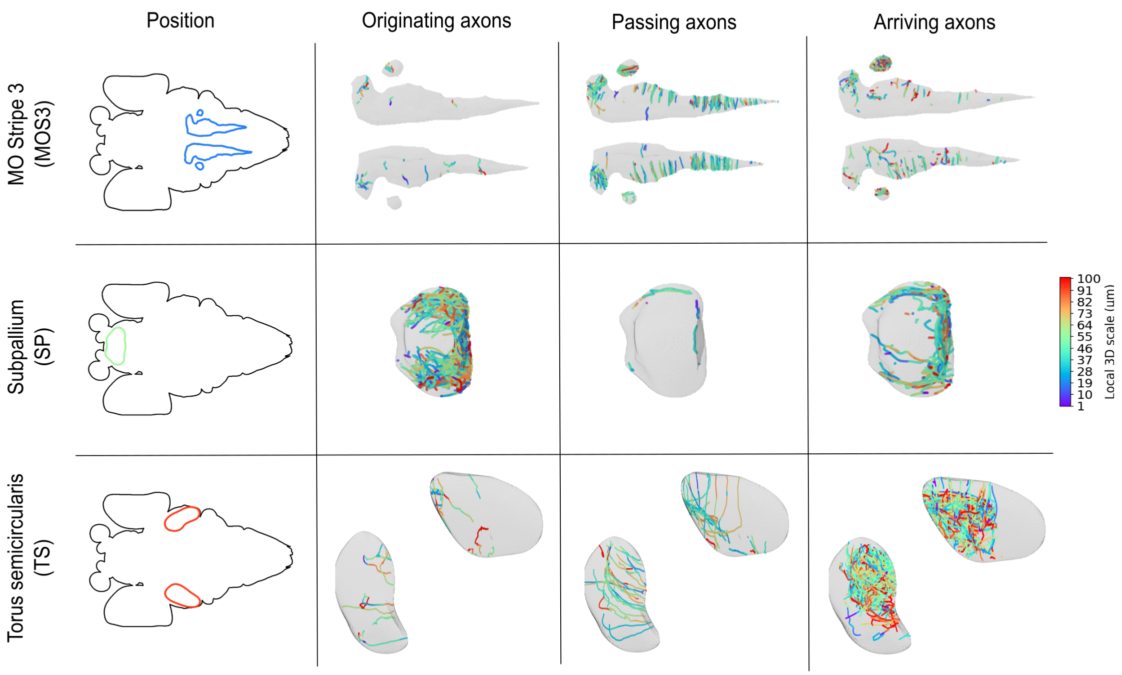


**Figure S11: Variability of local 3D scales in some brain regions (MOS3, SP, TS) for different connectivity axonal patterns (originating, passing, arriving).** The regional local 3D scale differs between originating, passing and arriving axons.

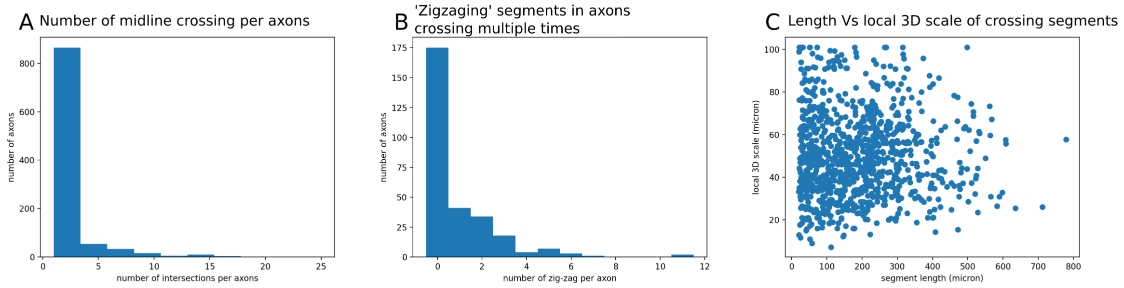


**Figure S12: Local 3D scale of axons crossing the midline.** (A) Distribution of the number of midline crosses made by individual axon arbors. (B) Distribution of the number of 'zigzags' through the midline, i.e. total number of crossings minus the number of crossing branches, for axons crossing more that 2 times. (C) Relation between the local 3D scale and length of crossing branches at the midline. A branch is defined as a neurite segment located between two branching point, or a branching point and a leaf or the root of the arbor.

**Table S1: Summary of 36 annotated brain regions from Kunst *et al.* (2019).**

| Index | Name | Abbr | Total_Nbaxon | Nbaxon_Orig | Nbaxon_Pass | Nbaxon_Arr | Mean_Local_3D_Scale |
|---|---|---|---|---|---|---|---|
| 1 | Medial_octavolateral_nucleus | MON | 281 | 115 | 20 | 146 | 50,81 |
| 2 | Cerebellum | Cb | 365 | 224 | 10 | 131 | 53,48 |
| 3 | MO_stripe_1 | MOS1 | 355 | 181 | 108 | 66 | 44,14 |
| 4 | MO_stripe_2 | MOS2 | 28 | 7 | 19 | 2 | 39,26 |
| 5 | MO_stripe_3 | MOS3 | 386 | 16 | 294 | 76 | 47,66 |
| 6 | MO_stripe_4 | MOS4 | 201 | 13 | 108 | 80 | 52,31 |
| 7 | MO_stripe_5 | MOS5 | 278 | 73 | 103 | 102 | 51,16 |
| 8 | interpeduncular_nucleus | IPN | 150 | 0 | 33 | 117 | 55,38 |
| 9 | inferior_olive | IO | 13 | 0 | 7 | 6 | 61,06 |
| 10 | caudal_hypothalamus | Hc | 70 | 2 | 10 | 58 | 48,56 |
| 11 | Raphe_nucleus | Ra | 332 | 8 | 140 | 184 | 55,28 |
| 12 | Tegmentum | T | 505 | 71 | 173 | 261 | 53,1 |
| 13 | anterior_reticular_formation | aRF | 727 | 16 | 216 | 495 | 54,41 |
| 14 | intermediate_reticular_formation | imRF | 581 | 18 | 214 | 349 | 52,33 |
| 15 | posterior_reticular_formation | pRF | 562 | 37 | 171 | 354 | 51,23 |
| 16 | Glossopharyngeal_ganglion | GG | 2 | 1 | 0 | 1 | 66,08 |
| 17 | Habenula | Hb | 34 | 6 | 3 | 25 | 51,71 |
| 18 | intermediate_hypothalamus | Hi | 366 | 21 | 47 | 298 | 55,38 |
| 19 | rostral_hypothalamus | Hr | 31 | 2 | 3 | 26 | 59,67 |
| 20 | Octaval_ganglion | OG | 19 | 3 | 10 | 6 | 47,8 |
| 21 | Olfactory_bulb | OB | 41 | 9 | 3 | 29 | 49,1 |
| 22 | Olfactory_epithelium | OE | 14 | 7 | 1 | 6 | 48,41 |
| 23 | Pallium | P | 110 | 18 | 6 | 86 | 51,13 |
| 24 | Pituitary | Pi | 5 | 0 | 0 | 5 | 67,33 |
| 25 | Posterior_teberculum | PT | 76 | 12 | 30 | 34 | 46,71 |
| 26 | preoptic_region | PO | 127 | 19 | 50 | 58 | 49,71 |
| 27 | pretectum | PrT | 231 | 68 | 55 | 108 | 54,65 |
| 28 | Retina | R | 51 | 47 | 1 | 3 | 60,71 |
| 29 | subpallium | SP | 105 | 67 | 3 | 35 | 52,94 |
| 30 | tectum | TeO | 195 | 57 | 2 | 136 | 54,2 |
| 31 | Thalamus | Th | 437 | 67 | 106 | 264 | 53,59 |
| 32 | Torus_longitudinalis | TL | 13 | 8 | 1 | 4 | 46,27 |
| 33 | Torus_semicircularis | TS | 195 | 11 | 50 | 134 | 57,53 |
| 34 | Trigeminal_ganglion | TG | 4 | 3 | 1 | 0 | 57,1 |
| 35 | Vagal_region | VR | 31 | 10 | 8 | 13 | 57 |
| 36 | vagus_motor_neurons | NX | 46 | 30 | 4 | 12 | 53,18 |

**Table S2: List of regions having axons originating from and arriving to the Torus Semicircularis (TS) from Kunst *et al.* (2019).** Index 0 corresponds to axons not starting from any regions.

| Index | Name | Abbr | Number |
|---|---|---|---|
| 0 | NaN | NaN | 43 |
| 1 | Medial_octavolateral_nucleus | MON | 21 |
| 2 | Cerebellum | Cb | 4 |
| 5 | MO_stripe_3 | MOS3 | 3 |
| 6 | MO_stripe_4 | MOS4 | 2 |
| 7 | MO_stripe_5 | MOS5 | 11 |
| 12 | Tegmentum | T | 6 |
| 13 | anterior_reticular_formation | aRF | 2 |
| 14 | intermediate_reticular_formation | imRF | 3 |
| 18 | intermediate_hypothalamus | Hi | 2 |
| 19 | rostral_hypothalamus | Hr | 1 |
| 25 | Posterior_teberculum | PT | 1 |
| 27 | pretectum | PrT | 5 |
| 30 | tectum | TeO | 28 |
| 31 | Thalamus | Th | 7 |
| 35 | Vagal_region | VR | 2 |
